# Evolution of reduced mate harming tendency of males in *Drosophila melanogaster* populations selected for faster life history

**DOI:** 10.1101/2021.10.30.466596

**Authors:** Tanya Verma, Anuska Mohapatra, Harish Kumar Senapati, Rakesh Kumar Muni, Purbasha Dasgupta, Bodhisatta Nandy

## Abstract

Detrimental effect of males on female, often termed mate harm, is a hallmark of sexual conflict. Allowed to evolve unchecked, mate harming traits are predicted to bring down average fitness of a population, unless mitigated by the evolution of resistance in females. In addition, life history may also modulate sexual conflict, but the mechanism is not clearly understood. Here we investigated the evolution of mate harm in a set of experimentally evolved laboratory populations of *Drosophila melanogaster* wherein a faster aging has evolved in response to >1000 generations of selection for faster development and early reproduction. We quantified mortality and fecundity of Oregon R females held with evolved (ACO) and ancestral males (CO) to show that the evolved males are significantly less detrimental to their mates. We compared our results from the ACO males with that from a phenocopied version of the ancestral regime (CCO) to show that only part of the observed difference in mate harm can be attributed to the evolved difference in body size. We further show that the reduction in mate harming ability evolved despite an increase in courtship activity, especially early in life. We discuss the causative role of an evolved reproductive schedule and altered breeding ecology.

**Significance statement:** Sexually antagonistic male effects can significantly bring down female fitness. Along with female counter evolution of resistance traits, life history has been conjectured to impose constrains on the evolution of such harming ability in males. Here, we report the evolution of mate harming ability in males of a set of five replicate *Drosophila melanogaster* populations that evolved smaller size and faster aging as a result of >1000 generations of experimental evolution for faster development and early reproduction. We show that in spite of ample scope of sexual selection, the faster aging males have evolved reduced mate harming ability despite being more active in courting their mates. To the best of our knowledge, this is one of the first clear evidences demonstrating the causal relationship between evolution of life history and reduction in sexual antagonism in a population.

## Introduction

Male fitness depends on traits that maximize mating and/or fertilization success. Such traits often includes coercive or manipulative traits that induces females to mate and/or reproduce at a rate that may benefit the males expressing them, even if they have detrimental side-effects to their mates (Parker 1979; Johnstone and Keller 2000; Chapman et al. 2003; Morrow et al. 2003; Arnqvist and Rowe 2005; Parker 2006; Queller and Strassmann 2018). While a locus that expresses such male benefiting trait evolves due to male specific selection, female specific selection, on the other hand, leads to counter-adaptation at a second locus that expresses female specific traits that allow females to either bypass or resist males coercion (Wigby and Chapman 2004; Nandy et al. 2013a; Dougherty et al. 2017; Chapman 2018; Rostant et al. 2020). Such evolutionary conflict between male trait loci and female trait loci is commonly referred to as interlocus sexual conflict (Chapman et al. 2003). Herein, both sexes are selected to evolve sex specific traits often resulting in antagonistic co-evolution between the sexes (Parker 1979; Rice 1992; Civetta and Singh 1995; Rice 1996; Arnqvist and Rowe 2002, 2005; Rönn et al. 2007). As interlocus sexual conflict continues, it may result in the evolution of male competitive traits such as, persistent courtship, extravagant display, traumatic insemination, deception, and seminal fluid proteins (Arnqvist and Rowe 2005, Koene 2012). For example, *Drosophila melanogaster* males overwhelm the females with incessant courtship, which often involve chasing, attempted mounting and other forms of physical interactions (von Schilcher and Dow 1977) that negatively impact female survival. Further, a copulating male transfers a complex cocktail of peptides/proteins (seminal fluid proteins), which increase egg production in females and make them less receptive to future mating (Wolfner 1997). The side-effects of these physiological alterations by these seminal fluid proteins is increased mortality (Chapman et al. 1995), and may even result in significant reduction in lifetime progeny output (Stewart et al. 2005). This detrimental effect of the males on their mates is often termed mate harm (Jiang et al. 2011, Nandy et al. 2013a, b, MacPherson et al. 2018).

In absence of female counter adaptation, the negative fitness impact of mate harm can potentially lead to extinction of a population/species, an outcome dubbed as the “tragedy of commons” (Le Galliard et al. 2005; Rankin and Kokko 2006; Rankin et al. 2007). However, such an extreme outcome of interlocus sexual conflict is unusual, because of, at least, three reasons. First, as mentioned earlier, female counter adaptation to mate harm can result in a chase-away process where evolution of mate harming ability of males is neutralised by the emergence of female resistance (Holland and Rice 1998, Holland and Rice 1999; Wigby and Chapman 2004; Friberg 2005; Rankin et al. 2011; Dougherty et al. 2017; Snow et al. 2019). Secondly, traits that inflict mate harm, i.e., the sexually antagonistic male traits, are energetically expensive to express and hence, increase in such traits is constrained by trade-offs involving costly life history traits (Wedell et al. 2006; Bonduriansky et al. 2008; Adler and Bonduriansky 2014; Lemaître et al. 2020). Thirdly, survivorship pattern and scheduling of reproduction can also put additional constraints on the evolution of mate harm, for example by restricting mating system and breeding ecology (Mital et al. 2021, 2022). The latter two theories are a more recent development in the field and we focus on them in the present investigation.

Contrary to a long standing perception, males pay a non-trivial reproductive cost (Partridge and Farquhar 1981, Kotiaho and Simmons 2003, Lane et al. 2010). If a male invests greater proportion of the finite amount of available resources in such reproductive traits, the potential to invest on other traits such as, soma maintenance and stress resistance physiology, immunity etc., is expected to reduce. This results in a negative correlation between investment in reproductive traits and these traits defining the cost of reproduction for males. Competitive traits including behaviour such as, courtship and copulation (Cordts and Partridge 1996, Clutton-Brock and Langley 1997, Bretman et al. 2013) and physiological traits such as, synthesis of a functional ejaculate (Dewsbury 1982; Andersson 1994; Arnqvist and Rowe 2005) are energetically expensive. Hence, male competitive traits constitute a significant part of the total reproductive cost. A comparative study on mammals indicated a significant association between degree of sexual selection on males and male biased reproductive cost (Promislow 1992). Experimental relaxation of sexual selection by enforced monogamy led to reduced investment in such costly competitive traits, including ejaculate composition, in *D. melanogaster* males and incidentally turned such males into less harming mates (Pitnick et al. 2001). Trade-off between competitive traits and somatic maintenance is a major contributor to the reproductive cost in males (Bonduriansky et al. 2008). Therefore, investment in longevity traits is expected to constrain the expression of male reproductive traits (Partridge and Farquhar 1981; Stearns 1989; Cordts and Partridge 1996; Clutton-Brock and Langley 1997). Life history theory predicts that compared to populations with shorter lifespan, males in populations with longer lifespan (i.e., higher investment in somatic maintenance) may have limited potential to invest in reproductive traits (Rose and Charlesworth 1981; Ernsting and Isaaks 1991; Kirkwood and Rose 1991; Service 1993; Tatar et al. 1993; Kotiaho 2001; Hunt et al. 2004). Though it is not clear whether lifespan evolution is caused by changes in reproductive traits or the other way round or both. Nonetheless, on an average, populations with shorter male lifespan are expected to have the potential to invest more in reproductive traits, including competitive traits involved in inflicting incidental mate harm, especially when there is ample selection for increased male competitive ability. Interestingly, existing evidences suggest that variation in lifespan and aging rate among males appears to reflect variation in reproductive investment (Alcock 1996; Cordts and Partridge 1996; Clutton-Brock and Langley 1997; Prowse and Partridge 1997; Hunt et al. 2004; Bonduriansky and Brassil 2005). Whether such correlation can also be extended to include sexually antagonistic male traits is not clear.

Scheduling of reproduction along with adult lifespan can directly modulate the extent of sexual conflict by constraining the breeding ecology. For example, a semelparous species with only one breeding season may tend to be monogamous and hence, experience reduced sexual conflict compared to an iteroparous species (Montrose et al. 2004; Clutton-Brock 2017; Griffith 2019). Long et al. (2010) showed that in a set of *Drosophila* laboratory populations, which are effectively semelparous, timing of mating from the semelparous breeding window, could significantly alter the sexually antagonistic outcomes in females. Bonduriansky (2014) showed that change in background mortality rate, resulting from increased predation pressure, could result in a short-term relaxation of sexual conflict. Population with low female life expectancy, increased mate harm by males is expected to bring down the average fitness of the population (Rankin and Kokko 2006) creating a scenario where selection can act against mate harming traits. If shorter lifespan evolves as a result of selection of components of life history (for example, faster pre-adult development and maturation), it can significantly shorten the potential duration of male-female encounter thereby reducing sexual selection and conflict in a population (Milat et al. 2021, 2022). There are several clear evidences supporting the notion that male-female encounter rate is, indeed, an important determinant of variation in interlocus sexual conflict across populations. For example, change in (a) the complexity of the physical environment (Byrne et al. 2008; Yun et al. 2017), (b) thermal environment that alters various activities in males (García-Roa et al. 2019), and (c) community structure that alters male-female encounter rate (Gomez-Llano et al. 2018) have been found to alter the level of mate harm in a population.

Here, we investigated the evolution of sexually antagonistic male traits in a set of experimentally evolved populations of *D. melanogaster* having substantially faster pre-adult development and reduced lifespan compared to their ancestors due to selection for faster development and early reproduction for over 1000 generations. These selected populations – ACO (Accelerated CO) will be hereafter referred as ‘faster aging population’, and their ancestral populations – CO (Control derived from O’s, which themselves were a set of populations selected for reproduction at an Old age) as ‘ancestral/control population’. The details of the history of these populations can be found in Chippindale et al. (1997). The faster aging populations have been extensively investigated for a range of life history traits since early 1990’s. As a direct response to selection, ACO populations evolved ~20% reduction in pre-adult development time and >18% reduction in mean life expectancy (Chippindale et al. 1997). Moreover, the maintenance regime of these populations is such that their effective adult lifespan is quite dramatically short (see Methods). In effect, ACO life history has evolved to adopt the so-called “live fast-die young” life history. Interestingly, in their short adult life, there is a substantial scope for scramble competition for mating. However, selection under such an ecology, can be expected to operate on variation that affects early-life fitness components, including competitive ability and courtship performance. Given that investment in the physiology of somatic maintenance has significantly reduced (as a result of shorter lifespan), theories would, thus, predict a resource re-allocation whereby male early-life competitive traits are preferentially allocated. However, apart from significantly reduced life expectancy, ACO flies are also visibly smaller in size – possibly due to the effect of selection for faster development. Therefore, compared to the controls, these faster aging populations also have significantly reduced total resource that is available for allocation in different physiological processes, including those connected to the expression of competitive traits. Thus, pertaining to the evolution of mate harming ability of ACO males, we have two contrasting predictions.

To test the above mentioned theories, we first quantified mate harm inflicted by the control and evolved males on a standard female type. Then, we specifically compared mating behaviour of these males to assess the divergence in the physical component of mate harassment. Apart from males from the evolved faster aging populations and their matched controls, we used a third treatment where experimental control males were phenocopied to match the size of the faster aging males. This third treatment was used to tease out the effects of genetically evolved difference in mate harm, and that cause solely due to reduced size. We used a standard laboratory line, Oregon R, as the common female background against which mate harming ability of the treatment males were compared. Importantly, we compared all the measured traits across the evolved and ancestral populations at five different age-points to assess the evolved difference in age-specific expression of the sexually antagonistic male traits.

## Methods

As mentioned above, the investigation reported here used ten laboratory populations of *Drosophila melanogaster* – five ACO populations and their matched ancestral CO populations. These populations were kindly provided to us by Prof. Michael R. Rose of University of California, Irvine, USA. They were maintained for 20 (CO) and 80 (ACO) generations in our laboratory before starting the experiment. The details of population history can be found in Chippindale et al. (1997). A brief description is provided in Figure S1 in the supplementary information. On November 1991, five replicate CO populations – CO1-5 (subscript refers to replicate identity), were used to initiate five replicate populations selected for accelerated development, viz. ACO1-5 (Rose et al. 1992). CO populations are maintained on a four-week discrete generation cycle, under 24-hours light, 25 °C (± 1), ~80% relative humidity on standard banana-jaggery-yeast medium. Larval density is controlled at ~70 per 8ml medium in each culture vial by culturing the desired number of eggs in growth vials. 40 such vials constitute a population. On day 12 following egg culture, adults from all 40 vials are transferred to a population cage. The CO flies take about 9-10 days to complete pre-adult development and therefore, by day12 virtually all surviving flies finish development and are in adult stage. A population cage is supplied with food on a petri dish. From day14 of the generation cycle, an old food plate is replaced with a fresh one every alternate day until day24. On day26, food plate smeared with *ad-lib* amount of yeast paste (paste made by dissolving baker’s yeast granules in water) is provided in the cage, replacing the old food. Approximately 48 hours following this, i.e., on day28, an oviposition substrate (two pieces of food cut in a trapezium shape) is introduced inside the cage, and a window of 18 hours is allowed for oviposition. Eggs deposited on the substrate, especially on the vertical slants, are collected by cutting out pieces of food with ~70 eggs in it. These pieces are introduced in fresh food vials to start the next generation.

The derived ACO populations employed a selection paradigm that involved strong selection for faster pre-adult development in addition to a much reduced, viz., approximately 24-36 hours of adult life (Chippindale et al. 1997). At the beginning of the selection lines, selection was imposed by allowing only fastest 20% developing flies to populate a generation, followed by brief window (~24-36 hours) of time for reproduction. Upon several hundreds of generations of selection, this maintenance was slightly modified. Their current maintenance regime involves a 9-day discrete generation cycle. The pre-adult development takes about 7-8 days. The adults are transferred to a population cage on day8 of the generation cycle. This cage is provided with standard food pieces presented as semi-circular oviposition substrate smeared with *ad-lib* amount of yeast paste. Eggs are collected in a manner similar to that used for the CO populations, following 24 hours of introducing the flies into the cage, to start the next generation, and ending the nine days long generation cycle. All other components of the ecology of the ACO populations are identical to those of the CO populations. The ACO populations had passed through >1200 generations by the time the following experiments were conducted.

### Generation of experimental flies

All experiments described here were done following the standard paradigm of experimental evolution assays in which both ACO and CO populations were passed through one generation of standard rearing, including a 14-day rearing schedule for CO’s and 13-day rearing schedule for ACO’s, to equalize the non-genetic parental effects. All experimental flies were reared at a density of 70 eggs per 8ml medium in a vial under the standard population maintenance regime described above. In all assays, to account for the difference in development time between ACO and CO flies (Chippindale et al. 1997), ACO eggs were collected two days following the collection of the CO eggs. This roughly synchronized the emergence of adult flies. Hereon, age of the adult flies refers to post-eclosion age, in days, unless mentioned otherwise.

We adopted an experimental design in which all male traits including the extent of sexually antagonistic effect of the experimental males was assessed against a common female background, which is phylogenetically unrelated to either male types, potentially equalizing the effect of female counter adaptation on the final outcome of male effect on female fitness. Further, using females from an inbred line, Oregon R, as a common background, we increased the resolution of assay as such inbred females are expected to be more susceptible to mate harm (Snook 2001, Pitnick and Garcia-Gonzalez 2002). This could maximise our ability to detect even relatively smaller difference in harming ability of males across the two regimes. Eggs were collected from Oregon R line and females were raised in the same manner as that followed for the CO and ACO males stated above, including the larval density of 70 per 8 ml food in a vial. All experimental females were age-matched with the experimental males.

Past selection has resulted in a considerable reduction in the body size of the ACO flies in both sexes (Passananti and Matos 2004; Burke et al. 2010, also see body weight data in the results section). Since smaller males have been previously shown to be less harming to their mates (Pitnick and García–González 2002; Friberg and Arnqvist 2003), the difference in body size between ACO and CO males was a confounding factor in our mate harm assay. We adopted a conservative strategy by introducing an additional treatment in our mate harm assay. In this treatment, we used smaller CO males having comparable body size as ACO males (hereafter referred to as phenocopied CO males, or CCO males). The phenocopied CO males were generated by growing them at the larval density of 240 per 3ml food in a standard vial, hence the name CCO (i.e., crowded CO). All the assays mentioned in later sections except courtship frequency and components of courtship behaviour were conducted using males belonging to three regimes – ACO, CO and CCO. The dry body weight at eclosion of the CCO males were not identical to that of the ACO males (see dry body weight results below). However, both ACO and CCO males were found to be smaller compared to the ancestral CO males by a comparable degree, approximately 50-54% lighter than the CO males. To put it simply, the evolved ACO males in our experiments were compared with the ancestral males as well as ancestral males which were phenocopied for smaller body size. We argue that if the difference in a trait between ACO and CO regimes is qualitatively identical to that observed between the CCO and CO regimes, the ACO-CO difference can be majorly attributed to the evolved body size difference. For example, if both ACO and CCO males induce reduced mate harm to the experimental females compared to that induced by the CO males, the evolved difference in the ACO males’ harming ability can be attributed to the smaller body size.

All the adult flies used in the experiments mentioned below were collected as virgins. At the onset of eclosion (emergence from pupal shell), virgin males and females were collected within 4-6 hours of eclosion. Flies were held in groups of 10 individuals per vial with ample food until the assay setup with alternate day food-change. A subset of the freshly eclosed males were frozen in −20°C for dry weight measurement. For every population we measured dry weight of 50 males. Frozen males were first air-dried in 60°C for 48 hours, before being weighed in a semi-microbalance in groups of five. All adult collections were done under light CO_2_ anaesthesia. The design and details of the individual assays are mentioned below. A generalized experimental plan is depicted in Fig.1.

**Fig.1.**
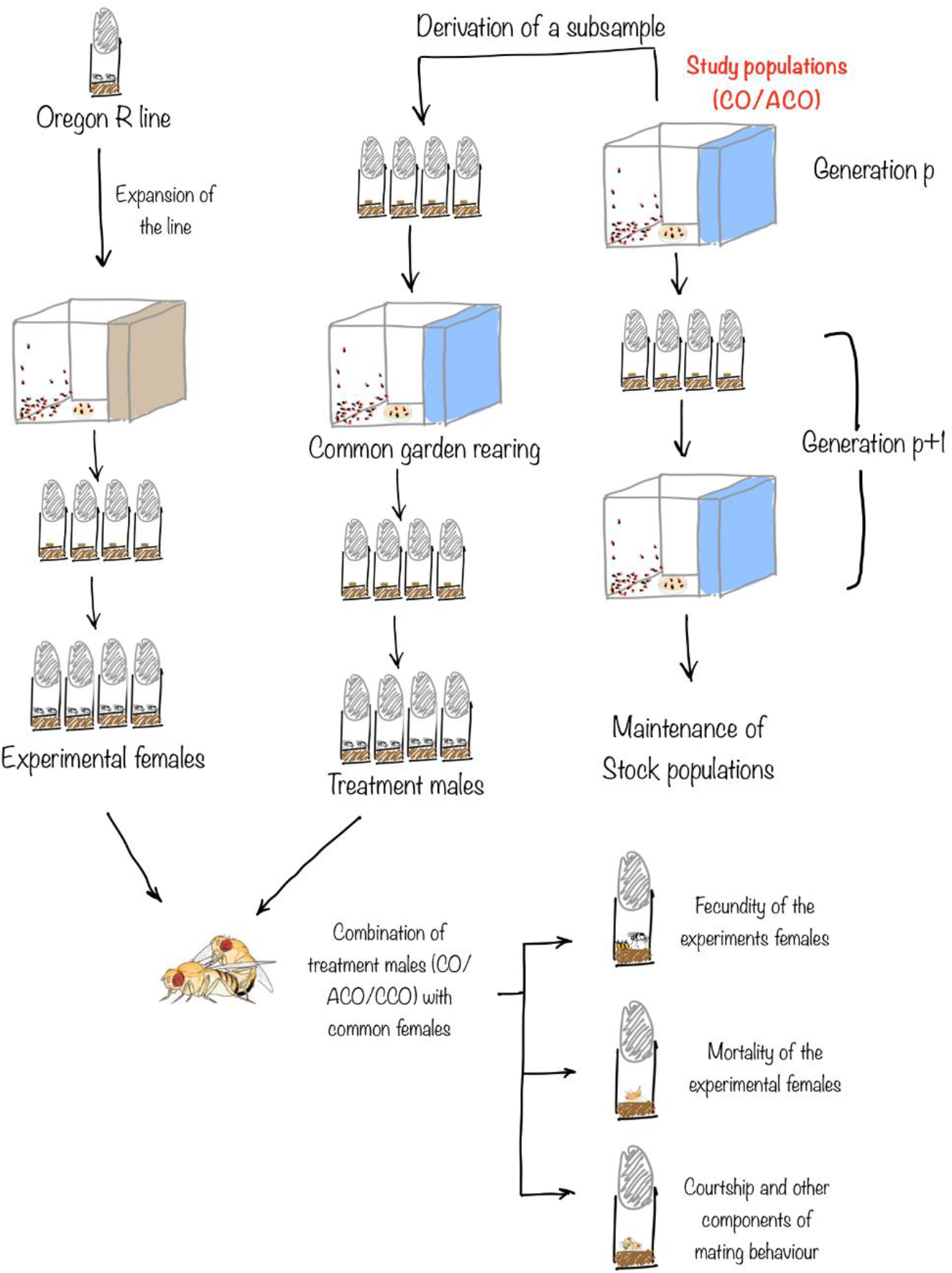
Schematic representation of the design main assay. Experimental subsets were generated from stock populations following one generation of standardization (common garden rearing). Experimental males were generated in the following manner: CO subset was used to generate CO experimental males and CCO males (phenocopied to ACO, these were generated by growing larvae at a density of 240/3ml of standard food) and ACO males were generated from ACO stock. Oregon R females were generated separately under standard conditions as common female for all three regime males. On assay day the whole set-up was divided into two male-exposure condition i.e. single exposure, SE (where males and females are allowed to mate once) and continuous exposure, CE (where males and females were housed together after first mating). Behaviour assays were then carried out for 20 days where female mortality was recorded every day, fecundity and courtship frequency was noted at every 5-day interval). All vials were flipped into fresh food vial at every alternate day

### Assay Setup

When adult flies were 1-2 days old, the assay vials were set up by combining males and females in food vials. Each of these vials, therefore, had ten males and ten females. A set of twenty vials was set up for a given population. The vials were left undisturbed for one hour and the flies were allowed to mate. Mating was visually observed. No video recording was done. Vials where all ten females did not successfully copulate were removed, leaving the final number of vials at this stage at 18-20 for each population. The entire set of these vials were divided into two male exposure assay conditions – (a) single mating (8-10 vials), and (b) continuous exposure (8-10 vials) treatments. In the single mating subset, after completion of the first round of mating, sexes were separated under light CO_2_ anaesthesia, and females were retained in the same vial while the males were discarded. The continuous exposure vials were retained without separating the sexes. Flies in these vials were also exposed to light CO_2_ anaesthesia to equalize the handling across the two subsets. All vials were maintained, with alternate day food-change, for twenty days. In these twenty days, mortality, fecundity and behaviour components were recorded (see below). Throughout the experiment, except sorting of sexes, all other fly handling including combination of sexes and transfer of flies from spent vials to fresh food vials were done without anaesthesia.

### Fecundity and mortality in test females

Female mortality was recorded daily for the all twenty days of the assay. Dead flies were removed from the vials by aspiration. Fecundity was recorded in approximately five day intervals, on day 1, 5, 10, 15 and 20. On each of these days, flies (only females for single mating and both sexes for continuous exposure set) from a vial were transferred to a fresh food vial (hereafter, fecundity vial) and allowed to lay eggs for a duration of ~24 hours after which they were transferred again into a fresh food vial to continue the assay (except day 20 count). The fecundity vials were then frozen immediately to stop further development. The eggs were later counted under a microscope. Per capita fecundity on a given assay day was calculated for a vial by dividing the total number of eggs in that vial by the number of females alive at the start of that day. Per capita fecundity values from individual vials were taken as the unit of analysis. For the analysis of the mortality data, proportion of females (i.e., out of a total ten in a vial) that were recorded dead at the end of the 20-day period in a vial, i.e., cumulative female mortality, was taken as the unit of analysis.

### Courtship frequency

Courtship frequency was measured as the average number of courtship bouts a male was found to perform per unit time. The continuous exposure vials were used for this purpose. Hence, courtship frequency of our treatment males was measured against Oregon R females. Since body size is not known to systematically affect courtship frequency, CCO males were excluded from this assay. Courtship measurement was done on day2, 6, 11, 16 and 20 of the assay. On a given observation day, four observations (n_c_ = 4) were recorded for a vial, maintaining a gap of 90 minutes between two subsequent observations. During an observation, a given vial was measured four times in quick succession in the following manner. A randomly picked male was observed for 15 seconds, during which the number of bouts of independent courtship (see later) events were counted. This was repeated four times for a vial, male selection being random each time. These four counts constituted one observation for that vial. An average for one such observation was calculated by dividing the total count by four, yielding an observation value (C_i_, where i = 1, 2, 3, 4). Mean courtship frequency was calculated for a replicate vial using the following function:

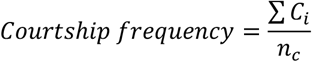

The observations were manual and did not involve any video recording. As the ACO and CO flies are visibly different, blind-folded observation did not have any utility and hence, was not adopted. Observers were well-trained to spot any of the following components of courtship behaviour: oriented toward female, following/chasing female, wing vibration, genitalia licking and attempted copulation (von Schilcher and Dow 1977). Performance of any of these behavioural components is counted as one bout of courtship as long as they were performed contiguously. During the observation, ‘two independent bouts of courtship’ is defined as (a) courtship events to two different females by the focal male, or (b) two courtship events performed by the focal male to the same female separated by either the male courting a different female or showing some other behaviour in between.

### Pattern of courtship behaviour in ACO and CO males

To investigate the qualitative difference in courtship behaviour ACO and CO regimes, we observed and quantified different components of courtship ritual in males in a separate trial. Courtship behaviour in male *D. melanogaster* is characterised by a complex series of discrete courtship components. In this assay, we quantified the frequencies of the five discrete components of courtship – (1) oriented toward female, (2) following/chasing female, (3) wing vibration, (4) genitalia licking, (5) attempted copulation (Ruedi and Hughes 2008). In addition to these five courtship components, males usually show three additional behavioural states – motionless, randomly moving, and copulation, which are not part of the courtship ritual (referred here as non-components). The observation vials were set up by introducing a 1-2 day old virgin male (either ACO or CO) and a 3-4 day old virgin Oregon R female in a fresh food vial without using anaesthesia. We started courtship observation after approximately 90 minutes from the initial introduction of the pair to the observation vial. This duration is usually sufficient for all females to undergo a single copulation (Nandy and Prasad 2011, Nandy et al. 2012, 2016). During observation, the behavioural state of the male in an observation vial was recorded by instantaneous scans. A given male was observed every 30 seconds for 30 minutes, resulting in a total of 60 observations for a male. This assay was done for all ten populations, i.e., all five replicate population pairs (ACO1-5 and corresponding CO replicate populations). In each population, 60 males were observed. Multiple trained observers (authors and volunteers) carried out the observations manually. However, for a given male all 60 observations were carried out by a single observer. Vials were randomly assigned among the observers to minimize observer bias in the data. Frequency of a component (for example, oriented toward female) for a male was calculated by using the following definition:

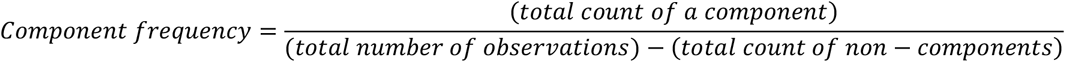

Thus, the assay resulted in 60 component frequency values (corresponding to the 60 males assayed for a population) for each of the five courtship components, for a population. These values were used as the unit of analysis.

### Statistical analysis

Cumulative female mortality data were analysed using three-factor mixed model Analysis of Variance (ANOVA) where Male regime (levels: CO, ACO and CCO) and male exposure type (levels: single mating and continuous exposure) were modelled as fixed factors, and block (level: 1-5) was modelled as a random factor. Dry body weight, and (courtship) component frequency were analysed using two-factor mixed model ANOVA, with Male regime and block modelled as fixed and random factors respectively. As female mortality and component frequency data were calculated as proportion values, analysis was performed following arcsine square root transformation (Sokal and Rohlf 1995; Zar 1999). All multiple comparisons were done using Tukey’s Honestly Significant Difference (HSD). All these analyses were done using Statistica (Tibco Software Inc., version 13.3).

Fecundity (i.e., per capita fecundity) data was analysed in two ways. First, per capita fecundity pooled across the five age classes, i.e., cumulative fecundity was analysed using three-factor mixed model ANOVA with male regime and male exposure type were modelled as fixed factors, and block was treated as a random factor. Secondly, to assess the effect of continued exposure to males on female fecundity, age-specific per capita fecundity was analysed only for the continuous exposure set. Initial analysis indicated significant block-to-block variation (see Results section and supplementary information, Table S3). Hence, each block was separately analysed using a linear mixed-effect model in R version 3.6.1 using lme4 package (R Development Core Team, 2019) (Bates et al. 2014) and lmerTest (Kuznetsova et al. 2017). In the model, per capita fecundity was modelled as response variable; male regime, age, and their two-way interactions as fixed factors; Vial id (replicate vial identity) as a random factor. The model used type III sum of squares. Following model was used for analysis:

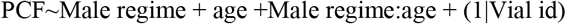

Courtship frequency was also analysed using `lmerTest` and `lme4` package in R. Courtship frequency was modelled as response variable with male regime, age and their two-way interaction as fixed factors and block including all its interactions as random factor. Vial id (replicate vial identity) nested within block was fitted as a random factor. Analysis indicated that block and its interactions were not significant contributors to the overall variance. Detailed analysis with random effects is provided in supplementary information (Table S6). The following linear-mixed model was used to analyse courtship frequency, while post-hoc comparisons were done using Tukey’s HSD using `emmeans` package (Lenth 2018):

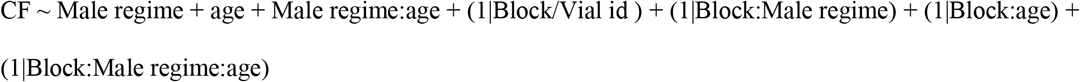

## Results

### Female mortality

Compared to the continuous exposure condition, where sexes were held together for the entire twenty days of the assay, female mortality was substantially lower under single mating assay condition (cumulative mortality <10%). Moreover, female mortality was substantially less when they were held with faster aging ACO males or the phenocopied CCO’s. Both the fixed factors, i.e., male regime and male exposure type, were found to have significant effects on cumulative female mortality (Table 1). In addition, the effect of the male regime × male exposure type interaction was also found to be significant (Table 1). Tukey’s HSD indicated that under single mating assay condition, the differences in cumulative female mortality across the three male regimes were not statistically significant. Under CE condition, however, it was substantially higher, especially when the females were held with CO males (>24% higher compared to either ACO or CCO males, Fig. 2a). The difference between ACO and CCO treatments was not significant. As the analysis indicated a significant effect of male regime × male-exposure type × block three-way interaction (Table 1), the results from each block were analysed separately using two-factor ANOVA (male regime and male-exposure type as fixed factors). The detailed outcome of these analyses can be found in the supplementary material (Table S5). Briefly, block 3 failed to show any effect of the regime, results from all other blocks were qualitatively identical.

**Table 1.** Summary of results of three-factor ANOVA on cumulative female mortality, cumulative fecundity, and two-factor ANOVA on dry body weight of males at eclosion. Mortality analysis was done on arcsine square root transformed values. Regime and male-exposure type are taken as fixed factor and block as random factor in female mortality and cumulative fecundity analysis. Regime is taken as fixed factor and block as random factor in dry body weight analysis. All tests were done considering α=0.05 and significant p-values are mentioned in bold font style. Abbreviations, MR: Male regime; MET: Male exposure type

| Trait | Effect | SS | DF | MS | Den DF | Den MS | F | p |
| --- | --- | --- | --- | --- | --- | --- | --- | --- |
| Female mortality | MR | 2.580 | 2 | 1.290 | 8.005 | 0.080 | 16.075 | **0.001** |
|  | MET | 6.933 | 1 | 6.933 | 4.002 | 0.070 | 98.434 | **<0.001** |
|  | Block | 0.831 | 4 | 0.208 | 0.371 | 0.037 | 5.636 | 0.550 |
| | MR × MET | 4.560 | 2 | 2.280 | 8.004 | 0.114 | 20.032 | **0.001** |
| | MR × Block | 0.642 | 8 | 0.080 | 8.000 | 0.114 | 0.705 | 0.684 |
| | MET × Block | 0.282 | 4 | 0.070 | 8.002 | 0.114 | 0.619 | 0.661 |
| | MR × MET × Block | 0.911 | 8 | 0.114 | 265.000 | 0.047 | 2.401 | **0.016** |
| Dry body weight | MR | 0.543 | 2 | 0.271 | 8.000 | 0.004 | 65.570 | **<0.001** |
|  | Block | 0.037 | 4 | 0.009 | 8.000 | 0.004 | 2.217 | 0.157 |
| | MR × Block | 0.033 | 8 | 0.004 | 135.000 | 0.001 | 6.781 | **<0.001** |
| Cumulative fecundity | MR | 206.30 | 2 | 103.150 | 8.001 | 12.371 | 8.338 | **0.011** |
|  | MET | 170.890 | 1 | 170.890 | 4.000 | 14.423 | 11.849 | **0.026** |
|  | Block | 1322.53 | 4 | 330.630 | 6.482 | 21.919 | 15.085 | **0.002** |
| | MR × MET | 104.850 | 2 | 52.430 | 8.002 | 4.877 | 10.750 | **0.005** |
| | MR × Block | 98.990 | 8 | 12.370 | 8.000 | 4.877 | 2.537 | 0.105 |
| | MR × Block | 57.690 | 4 | 14.420 | 8.002 | 4.877 | 2.958 | 0.090 |
| | MR × MET × Block | 39.020 | 8 | 4.880 | 267.000 | 3.044 | 1.602 | 0.124 |

**Fig. 2.**
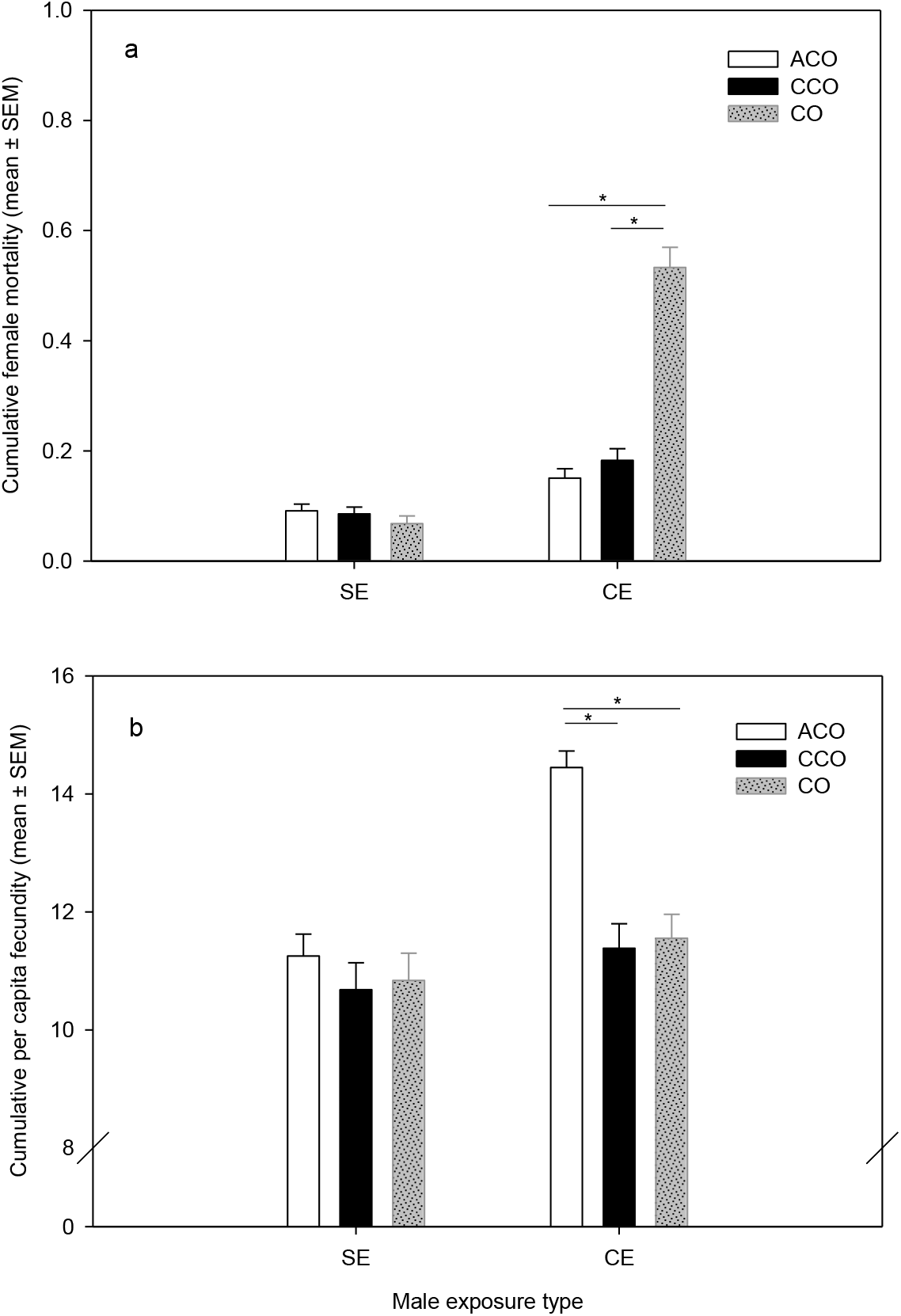
Effect of exposure to treatment males (ACO/CCO/CO) on female mortality and fecundity. (a) Proportion of females died by the end of the 20-day assay period (cumulative female mortality), under the two male-exposure conditions – single exposure (SE) and continuous exposure (CE). The vertical bars indicate the mean across all replicate populations. Error bars represent the standard errors of means (SEM); (b) Average per capita fecundity of experimental Oregon R females exposed to treatment males across all five age classes, under different male exposure types SE and CE. Means were calculated over five replicate populations. Only relevant multiple comparisons, which were done using Tukey’s HSD, are shown. Significant differences are marked with horizontal line and an asterix (*)

### Dry body weight

Dry body weight results confirmed that ACO males were substantially smaller compared to their ancestral CO males. Further, phenocopied CCO males qualitatively mimicked a similar size reduction of ACO males compared to that of the CO’s. We found significant effect of regime (p < 0.001, Table 1). Pairwise comparisons using Tukey’s HSD showed that both ACOs and CCOs are significantly smaller than COs (weight in mg, mean ± standard error of mean, ACO: 0.162 ±0.003; CO: 0.271 ±0.004; CCO: 0.131 ±0.005). Qualitatively the pattern of dry body weight was similar in all the blocks, but we found significant interaction of regime × block. Hence, summary table of each block is provided in supplementary information (Table S1, Fig. S2).

### Female fecundity

Fecundity of females held with faster aging ACO males was found to be significantly higher compared to both the ancestral CO males as well as those from the phenocopied CCO treatment. While the trend in cumulative fecundity across all age points was quite clean, age specific fecundity results were more complex. Results of the three-factor mixed model ANOVA revealed significant effects of male regime, male exposure type, and male regime × male exposure type interaction on cumulative per capita fecundity (Table 1). Multiple comparisons using Tukey’s HSD indicated that under single mating exposure, the three regime did not differ significantly from each other. Under the continuous exposure assay condition, females held with ACO males were found to have ~13.65% higher cumulative per capita fecundity compared to the females held with CO males. The difference in cumulative fecundity of females held with CCO males and that of the females held with CO males was statistically insignificant (Fig.2b).

Results of the linear mixed model analysis on age-specific per capita fecundity of the females in the continuous exposure set were more complex (see supplementary information). While in each block significant effects of male regime, age and male regime × age interaction were detected, there was little consistent trend across blocks (Table S4). However, in at least three of the five age points, females held with ACO males showed a significantly higher per capita fecundity compared to the other two treatments, particularly at later age points (Fig. S3).

### Courtship frequency and pattern

The ACO males were found to be significantly more active in courting females. Linear mixed model analysis on the courtship frequency (i.e., CF) results showed significant effects of male regime, age, and regime × age interaction (Table 2). Tukey’s HSD results showed that courtship frequency of ACO males is significantly higher, particularly at early age classes than the CO males (Fig. 3a).

**Table 2.**
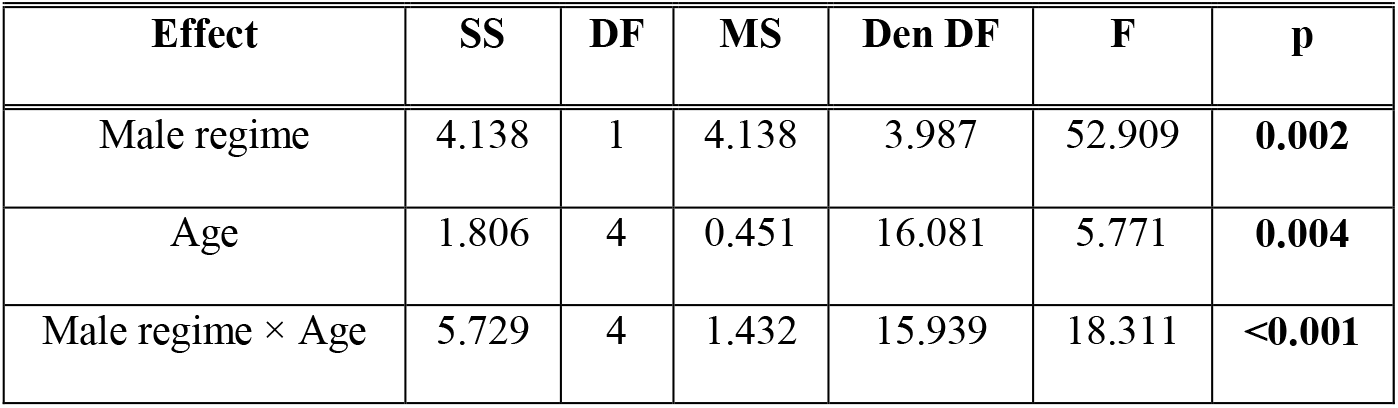
Summary of the results of linear mixed model (LMM) analysis of courtship frequency using `lmerTest` function in R. Regime and age were modelled as fixed factors and block as a random factor. All tests were done considering α=0.05 and significant p-values are mentioned in bold font style

| Effect | SS | DF | MS | Den DF | F | p |
| --- | --- | --- | --- | --- | --- | --- |
| Male regime | 4.138 | 1 | 4.138 | 3.987 | 52.909 | **0.002** |
| Age | 1.806 | 4 | 0.451 | 16.081 | 5.771 | **0.004** |
| Male regime × Age | 5.729 | 4 | 1.432 | 15.939 | 18.311 | **<0.001** |

**Fig. 3.**
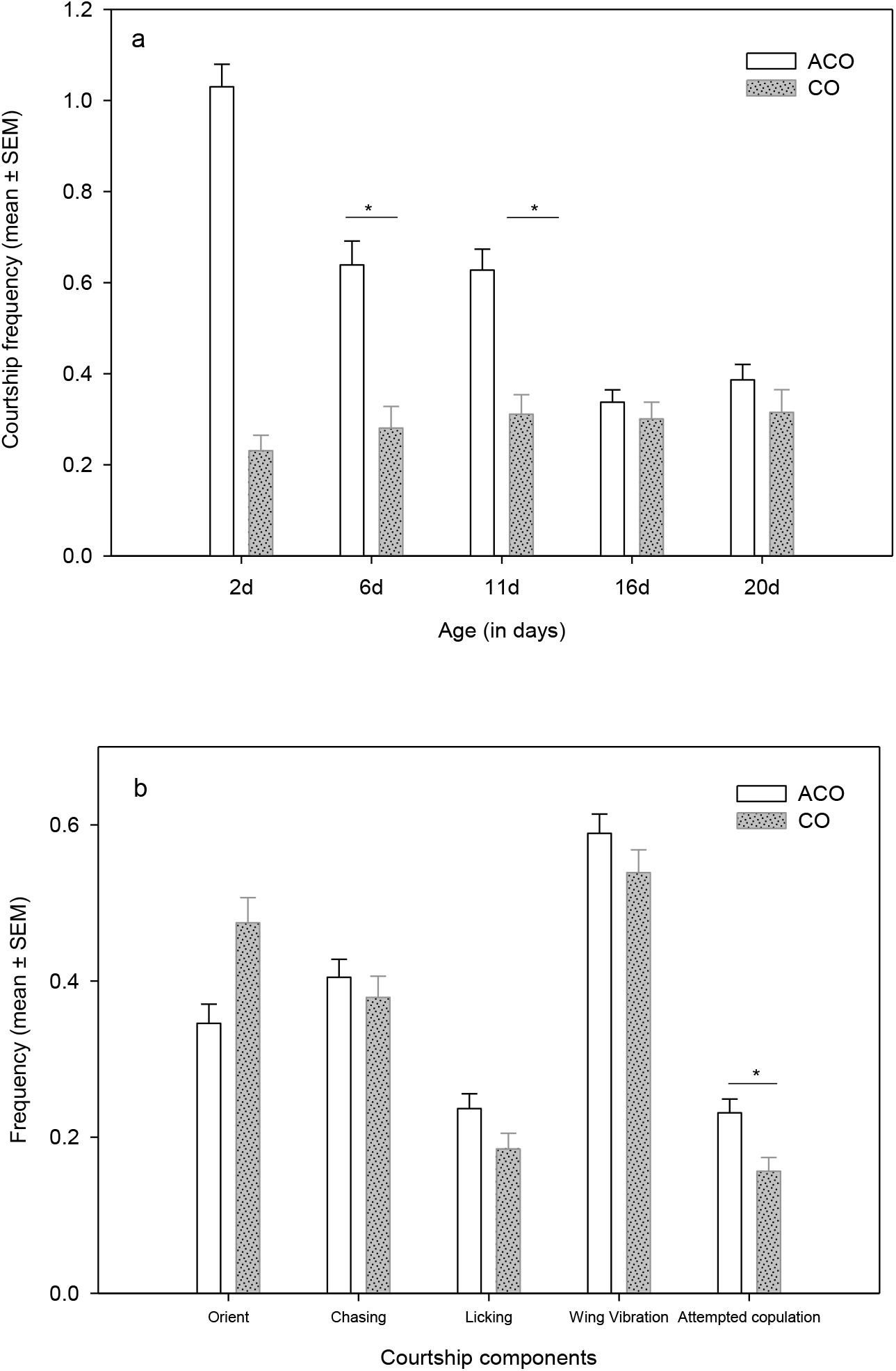
Results of the courtship behaviour assays. (a) Courtship frequency (bouts of courtship per observation) was measured for the 20-days assay period on days – 2, 6, 11, 16, and 20 in the CE vials of the main assay. (b) Frequency of the five components (proportional contribution of a courtship component) of courtship ritual were measured in a separate assay, where ACO and CO males were held with Oregon R females. The vertical bars indicate the mean across all five replicate populations. Error bars represent the standard errors of means (SEM). Only relevant multiple comparisons are shown in the figure. Significant differences are marked with horizontal line and an asterix (*)

When we compared the courtship ritual performed by the CO and ACO males, trying to assess if the courtship ritual has evolved in response to selection for accelerated life cycle, the latter showed a significantly higher mounting attempt, the final step in the courtship sequence. The results of two factor mixed model ANOVA performed on the frequency of different courtship components indicated an interesting effect of male regime. Male regime was not found to have a significant effect on four of the five courtship components (Table 3). However, we found a significant male regime effect on attempted copulation (p = 0.029), where ACOs showed ~15% higher copulation attempts compared to COs (Fig. 3b, Table 3).

**Table 3.**
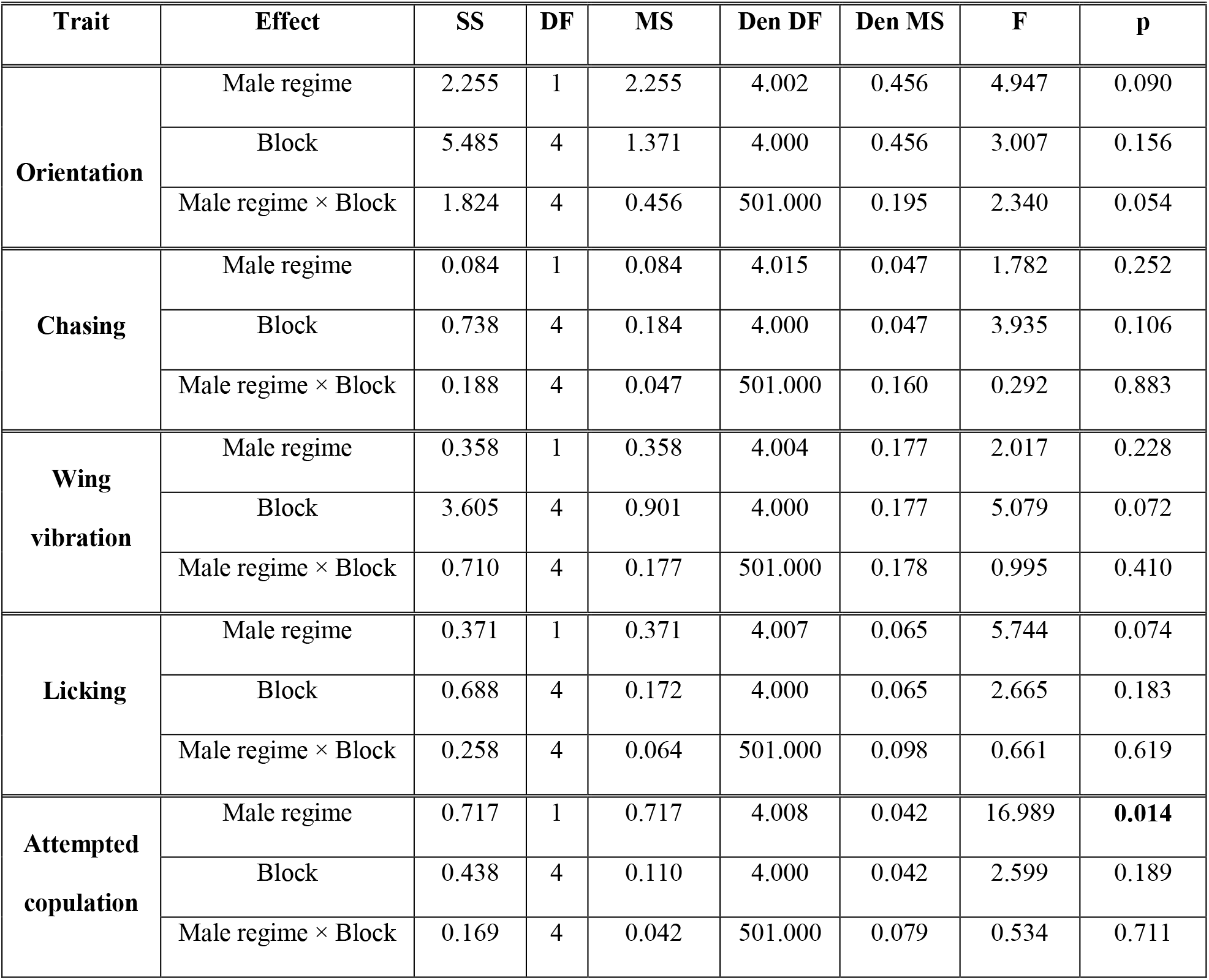
Summary of results of two-factor ANOVA on courtship component frequencies. Male regime was modelled as a fixed factor and block as a random factor. Analysis done on arcsine square root transformed values. All tests were done considering α=0.05 and significant p-values are mentioned in bold font style

| Trait | Effect | SS | DF | MS | Den DF | Den MS | F | p |
| --- | --- | --- | --- | --- | --- | --- | --- | --- |
| **Orientation** | Male regime | 2.255 | 1 | 2.255 | 4.002 | 0.456 | 4.947 | 0.090 |
|  | Block | 5.485 | 4 | 1.371 | 4.000 | 0.456 | 3.007 | 0.156 |
| | Male regime × Block | 1.824 | 4 | 0.456 | 501.000 | 0.195 | 2.340 | 0.054 |
| **Chasing** | Male regime | 0.084 | 1 | 0.084 | 4.015 | 0.047 | 1.782 | 0.252 |
|  | Block | 0.738 | 4 | 0.184 | 4.000 | 0.047 | 3.935 | 0.106 |
| | Male regime × Block | 0.188 | 4 | 0.047 | 501.000 | 0.160 | 0.292 | 0.883 |
| **Wing vibration** | Male regime | 0.358 | 1 | 0.358 | 4.004 | 0.177 | 2.017 | 0.228 |
|  | Block | 3.605 | 4 | 0.901 | 4.000 | 0.177 | 5.079 | 0.072 |
| | Male regime × Block | 0.710 | 4 | 0.177 | 501.000 | 0.178 | 0.995 | 0.410 |
| **Licking** | Male regime | 0.371 | 1 | 0.371 | 4.007 | 0.065 | 5.744 | 0.074 |
|  | Block | 0.688 | 4 | 0.172 | 4.000 | 0.065 | 2.665 | 0.183 |
| | Male regime × Block | 0.258 | 4 | 0.064 | 501.000 | 0.098 | 0.661 | 0.619 |
| **Attempted copulation** | Male regime | 0.717 | 1 | 0.717 | 4.008 | 0.042 | 16.989 | **0.014** |
|  | Block | 0.438 | 4 | 0.110 | 4.000 | 0.042 | 2.599 | 0.189 |
| | Male regime × Block | 0.169 | 4 | 0.042 | 501.000 | 0.079 | 0.534 | 0.711 |

## Discussion

Our results showed that, in spite of persistent harassment through increased courtship and copulation attempts, ACO males are clearly far less harming compared to the ancestral CO males. The females held with ACO males showed significantly less mortality and higher fecundity during the assay period. Interestingly, ACO males were found to be more active in courtship, especially early in life. This age-specific pattern of change in courtship activity is consistent with the theory of selection for early life fitness components in the ACO males. Despite several previous investigations linking courtship activity to the level of mate harm caused by *D. melanogaster* males, ACO males were found to be substantially less harming, in spite of having higher courtship activity.

Though faster aging ACO males are expected to have the potential to invest a greater proportion of their resources in competitive traits, potentially making them more harming to their mates, our results clearly show a contrasting trend. One potential explanation of this observation is that the ACO males are substantially smaller and are hence, less harming - measured in terms of the mates’ survival rate and reproductive output. Such an effect of body size on harming ability of the males in this system has been previously shown (Pitnick 1991, Pitnick and García–González 2002). Our results, however, suggest a more complex picture. Based on the ACO vs. CCO comparison, we could attribute only the mortality effect to the reduced size of the ACO males. However, the fecundity effect of the ACO males was quite unique. Average cumulative fecundity of experimental females held with the ACO males was found to be significantly higher than that with CO males. However, no significant difference was found between cumulative fecundity of females held with CO and CCO males. Thus, it is reasonable to deduce that the difference in reproductive output of the experimental females held with ACO males and those held with CO males was unlikely to be an outcome of the size difference between the males from the two regimes. Thus, either due to reduction is size or due to changes in traits unrelated to size, or both, ACO males are significantly less harming to their mates.

*D. melanogaster* males are known to physically coerce females through persistent courtship (Fowler and Partridge 1989). Multiple lines of evidence have also shown the correlation between the degree of mate harm and intensity of courtship behavior (Nandy et al. 2013b; MacPherson et al. 2018). For example, Nandy et al. (2013b) experimentally evolved a set of populations under male biased operational sex ratio that resulted in increased harming ability in males, along with increased courtship frequency. MacPherson et al. (2018) showed that females seemed to avoid mate harm when they were held in complex holding chambers with ample hiding opportunities, possibly by escaping direct exposure to persistent courtship. Different components of courtship behaviour have been found to have ample genetic variation both in natural and laboratory populations of *Drosophila* (Markow and Hanson 1981; Gromko 1987; Ritchie and Gleason 1995; Colegrave et al. 2000; Snook et al. 2005; Dai et al. 2008). There has been ample evidence suggesting evolvability of courtship behaviour in *D. melanogaster* (Bedhomme et al. 2008; Nandy et al. 2013b). However, how evolution of courtship behaviour can contribute to the evolution of mate harming ability of the males is not clear. The fact that ACO males, in our study, were found to be less harming despite being more active in courting females, is a direct challenge to the conventional wisdom that draws a one-on-one connection between courtship and physical component of mate harm. Not only ACO males courted females more frequently, they also attempted copulation more often - a component of courtship that appears to be more coercive. Hence, our results indicate that the relationship between courtship behaviour and mate harm is more complex than previously anticipated.

In addition to persistent courtship, seminal fluid proteins (Sfp) transferred to females during copulation have been shown to bring down female survival rate (Chapman et al. 1995; Wigby and Chapman 2005; Wigby et al. 2020). Changes in Sfp content of the ejaculate can potentially explain reduction in harming ability of the ACO males. At this point we do not have data on Sfp to directly test this hypothesis. However, in a separate assay, we observed that copulation between an ACO male and a female from the corresponding population is significantly shorter compared to that between a pair from a CO population (see SI for more details, Fig. S4 and Table S7, mean CD in minutes ±SEM, ACO: 16.15 ±0.657; CO: 20.32 ±0.623). Variation in copulation duration has been correlated with variation in the amount of Sfp transferred (Singh and Singh 2004; Friberg 2006; Bretman et al. 2009; Bretman et al. 2010). In principle, reduction in post-copulatory sexual selection can bring about changes in Sfp and/or copulatory traits in ACOs. Monogamy and female biased operational sex ratio have already been shown to lead to reduction in both harming ability and investment in certain attributes of seminal fluid content including Sfp (Holland and Rice 1999; Wigby and Chapman 2004; Crudgington et al. 2005; Linklater et al. 2007). However, it is unrealistic to suggest that our ACO populations have a monogamous breeding system and hence, post-copulatory sexual selection is absent. Our observations suggest that females in ACO populations undergo remating quite regularly (Fig. S5). However, the breeding ecology of this regime is vastly different from the ancestral regime. In contrast to the ancestral CO regime, where breeding life is approximately 19 days of adult life, ACO flies get 24-36 hours to reproduce after becoming adults. In this incredibly short breeding life, males are selected to be reproductively active and invest heavily on competitive traits that maximize mating success. Our courtship frequency results support this hypothesis. However, given that there is strong last male sperm precedence in this species (Manier et al. 2010) and ample female remating, the exact nature and intensity of post-copulatory sexual selection is not clear. A related possibility is that the experimental females held with ACO and CO males remated at different rates thereby received different quantities of Sfp. Though we do not have data on remating frequency in our assay, separate observation on these populations do not suggest difference in remating rate (see Fig. S5 in supplementary information).

In an independent investigation on a similar set of experimentally evolved populations (Ghosh and Joshi 2012; Mital et al. 2021, 2022) found that the intensity of sexual selection and the degree of interlocus sexual conflict to be lower in populations selected for faster development and early reproduction. The parallel between our results and those of Ghosh and Joshi (2012) and Mital et al. (2021, 2022) is somewhat expected, and is a robust proof in support of our theory. However, even more interesting is the difference. Whereas (Mital et al. 2021) could almost entirely assign the reduction in sexual antagonism in their faster developing and aging populations (viz., FEJ populations) to the reduction in body size, our results point to a different explanation. Comparison of the accelerated life cycle regime in Mital et al. (2021) and that of our ACO populations is interesting. In ACO regime, due to a fixed egg-to-adult development time window of eight days, selection for faster pre-adult development has not been strong for the last several hundred generations. However, the accelerated life cycle regime in Mital et al. (2021) has a strong directional selection for faster development, and as a result, flies under this regime have evolved a more extreme reduction in size, amounting to resource deprivation that did not allow the evolution of male competitive traits, including courtship (Mital et al. 2021). In addition, whereas the accelerated life cycle regime of Mital et al. (2021) practically ensures life-long monogamy, we observed substantial amount of promiscuity in ACO flies. Hence, it is possible that ACO males were specifically selected for traits that maximize mating and fertilization success in such a way that reduces incidental mate harm. Such results clearly show the importance of an incredibly nuanced breeding ecology of animals, and its connection to life history.

Theories have long predicted a connection between life history and sexual conflict. Most often, the trade-offs between costly sexually antagonistic traits and one or more life history trait(s), most notably somatic maintenance, has taken the centre stage in such discussions (Bonduriansky et al. 2008; Maklakov and Lummaa 2013; Hooper et al. 2017). However, if the aim is to understand and predict the evolutionary outcome of changes in life history on sexually antagonistic traits, perhaps a more important question is, how a given alteration in life history affects the breeding system, and by extension, the ecology of sexual selection. The secondary outcomes in that case can affect a wider range of sexually antagonistic traits, potentially independent of trade-offs. Our results on the reduced mate harming ability in ACO males is evidence in support of this thesis.

Rankin and Kokko (2006, 2007) very explicitly showed a seemingly obvious outcome of interlocus sexual conflict, i.e., population extinction due to unchecked evolution of sexually antagonistic male traits. However, most theories of interlocus conflict views such evolutionary dynamics as an arms’ race between sexes. Hence, counter adaptation in females is fundamental to the theory. Such counter adaptation is also expected to impede the evolutionary increase in the harming ability of males in a population, thereby avoiding the tragedy of commons. In line with such a reasoning, many empirical investigations have reported strong female counter adaptation to mate harm imposed by male (Wigby and Chapman 2004; Nandy et al. 2014). As we have argued above, life history can also be a potent constraint on the evolution of mate harm. However, there is little literature on this subject. To the best of our knowledge, the present study on the ACO males is one of the very few empirical evidence of evolutionary reduction in mate harming ability. The results presented here should be of general importance in explaining the variation in the intensity of sexual antagonism across species.

## Authors Contribution

Bodhisatta Nandy and Tanya Verma conceptualized the study, designed the experiments, and wrote the manuscript. Tanya Verma, Harish Kumar Senapati, Anuska Mohapatra, Rakesh Kumar Muni executed the experiments. Tanya Verma, Purbasha Dasgupta and Bodhisatta Nandy analysed and interpreted the results.

## Acknowledgements

The study was financially supported by a research grant from Department of Science and Technology, Govt. of India (INSPIRE Faculty award, Grant no. DST/INSPIRE/04/2013/000520). We thank Subhasish Halder for help in the experiments and data analysis. We thank Anish Koner and Rabisankar Pal for help in experimental observations. We thank Syed Zeeshan Ali for his valuable comments on a previous version of this manuscript and on the analyses. TV thanks Indian Institute of Science Education and Research, Berhampur for financial support in the form of Junior and Senior Research Fellowship. PD thanks Council for Scientific and Industrial Research, Government of India for financial support in the form of Junior and Senior Research Fellowship. AM thanks Department of Science and Technology, Govt. of India for financial support in the form of INSPIRE SHE scholarship.

## Date accessibility statement

A spreadsheet file with data from all assays has been included in the submission as a Supplementary material.

## References

Adler MI, Bonduriansky R (2011) The dissimilar costs of love and war: Age-specific mortality as a function of the operational sex ratio. J. Evol. Biol 24:1169–1177. https://doi.org/10.1111/j.1420-9101.2011.02250.x

Adler MI, Bonduriansky R (2014) Sexual conflict, life span, and aging. Cold Spring Harb. Perspect. Biol. 6:a017566. https://doi.org/10.1101/cshperspect.a017566

Alcock J (1996) Male size and survival: the effects of male combat and bird predation in Dawson’s burrowing bees, *Amegilla dawsoni*. Ecol. Entomol. 21:309–316. http://dx.doi.org/10.1046/j.1365-2311.1996.00007.x

Andersson M (1994) Sexual selection. Princeton University Press, Princeton, USA

Arnqvist G, Rowe L (2002) Antagonistic coevolution between the sexes in a group of insects. Nature 415:787–789. https://doi.org/10.1038/415787a

Arnqvist G, Rowe L (2005) Sexual conflict. Princeton University Press, Princeton, USA

Baer B, Maile R, Schmid-Hempel P, Morgan ED, Jones GR (2000) Chemistry of a mating plug in bumblebees. J. Chem. Ecol. 26:1869–1875. http://dx.doi.org/10.1023/A:1005596707591

Bates D, Mächler M, Bolker B, Walker S (2014) Fitting linear mixed-effects models using lme4. arXiv preprint arXiv:1406.5823

Bedhomme S, Prasad NG, Jiang P-P, Chippindale AK (2008) Reproductive behaviour evolves rapidly when intralocus sexual conflict is removed. PLoS One 3:2187. https://doi.org/10.1371/journal.pone.0002187

Bonduriansky R (2014) The ecology of sexual conflict: background mortality can modulate the effects of male manipulation on female fitness. Evolution 68:595–604. https://doi.org/10.1111/evo.12272

Bonduriansky R, Brassil C (2005) Reproductive ageing and sexual selection on male body size in a wild population of antler flies *(Protopiophila litigata)*. J. Evol. Biol 18:1332–1340. https://doi.org/10.1111/j.1420-9101.2005.00957.x

Bonduriansky R, Maklakov A, Zajitschek F, Brooks R (2008) Sexual selection, sexual conflict and the evolution of ageing and life span. Funct. Ecol 22:443–453. https://dx.doi.org/10.11012Fcshperspect.a017566

Bretman A, Fricke C, Chapman T (2009) Plastic responses of male Drosophila melanogaster to the level of sperm competition increase male reproductive fitness. Proc. Royal Soc. B 276:1705–1711. https://doi.org/10.1098/rspb.2008.1878

Bretman A, Fricke C, Hetherington P, Stone R, Chapman T (2010) Exposure to rivals and plastic responses to sperm competition in *Drosophila melanogaster*. Behav. Ecol 21:317–321. https://doi.org/10.1093/beheco/arp189

Bretman A, Westmancoat JD, Gage MJ, Chapman T (2013) Costs and benefits of lifetime exposure to mating rivals in male Drosophila melanogaster. Evolution 67:2413–22. https://doi.org/10.1111/evo.12125

Burke MK, Dunham JP, Shahrestani P, Thornton KR, Rose MR, Long AD (2010) Genome-wide analysis of a long-term evolution experiment with *Drosophila*. Nature 467:587–590. https://doi.org/10.1038/srep39281

Byrne PG, Rice GR, Rice WR (2008) Effect of a refuge from persistent male courtship in the *Drosophila* laboratory environment. Am. Zool 48:e1–e1. https://doi.org/10.1093/icb/icn001

Chapman T (2018) Sexual conflict: mechanisms and emerging themes in resistance biology. The Am. Zool. 192:217–229. https://doi.org/10.1086/698169

Chapman T, Arnqvist G, Bangham J, Rowe L (2003) Sexual conflict. Trends Ecol. Evol 18:41–47. https://doi.org/10.5860/choice.43-3391

Chapman T, Liddle LF, Kalb JM, Wolfner MF, Partridge L (1995) Cost of mating in *Drosophila melanogaster* females is mediated by male accessory gland products. Nature 373:241–244. https://doi.org/10.1038/373241a0

Chippindale AK, Alipaz JA, Chen HW, Rose MR (1997) Experimental evolution of accelerated development in *Drosophila*. 1. Developmental speed and larval survival. Evolution 51:1536–1551. https://doi.org/10.1111/j.1558-5646.1997.tb01477.x

Civetta A, Singh RS (1995) High divergence of reproductive tract proteins and their association with postzygotic reproductive isolation in *Drosophila melanogaster* and *Drosophila virilis* group species. J. Mol. Evol 41:1085–1095. https://doi.org/10.1007/bf00173190

Clutton-Brock T (2017) Reproductive competition and sexual selection. Philos. Trans. R. Soc. Lond. B, Biol. Sci 372:20160310. https://doi.org/10.1098/rstb.2016.0310

Clutton-Brock T, Langley P (1997) Persistent courtship reduces male and female longevity in captive tsetse flies *Glossina morsitans morsitans* Westwood (Diptera: *Glossinidae)*. Behav. Ecol 8:392–395. https://doi.org/10.1093/beheco/8.4.392

Colegrave N, Hollocher H, Hinton K, Ritchie M (2000) The courtship song of African *Drosophila melanogaster*. J. Evol. Biol 13:143–150. https://doi.org/10.1046/j.1420-9101.2000.00148.x

Cordts R, Partridge L (1996) Courtship reduces longevity of male *Drosophila melanogaster*. Anim. Behav 52:269–278. https://doi.org/10.1006/anbe.1996.0172

Crudgington HS, Beckerman AP, Brüstle L, Green K, Snook RR (2005) Experimental removal and elevation of sexual selection: does sexual selection generate manipulative males and resistant females? Am. Nat 165:S72–S87. http://dx.doi.org/10.1086/429353

Dai H, Chen Y, Chen S, Mao Q, Kennedy D, Landback P, Eyre-Walker A, Du W, Long M (2008) The evolution of courtship behaviors through the origination of a new gene in *Drosophila*. Proc. Natl. Acad. Sci. U.S.A. 105:7478–7483. https://doi.org/10.1073/pnas.0800693105

Dewsbury DA (1982) Ejaculate cost and male choice. Am. Nat. 119:601–610. https://doi.org/10.1086/283938

Dougherty LR, van Lieshout E, McNamara KB, Moschilla JA, Arnqvist G, Simmons LW (2017) Sexual conflict and correlated evolution between male persistence and female resistance traits in the seed beetle *Callosobruchus maculatus*. Proc. Royal Soc. B 284:20170132. https://doi.org/10.1098/rspb.2017.0132

Duxbury EM, Rostant WG, Chapman T (2017) Manipulation of feeding regime alters sexual dimorphism for lifespan and reduces sexual conflict in *Drosophila melanogaster*. Proc. Royal Soc. B 284:20170391. https://doi.org/10.1098/rspb.2017.0391

Ernsting G, Isaaks J (1991) Accelerated ageing: a cost of reproduction in the carabid beetle *Notiophilus biguttatus* F. Funct. Ecol 5:299–303. https://doi.org/10.2307/2389268

Fowler K, Partridge L (1989) A cost of mating in female fruit flies. Nature 338:760–761. https://doi.org/10.1038/338760a0

Friberg U (2005) Genetic variation in male and female reproductive characters associated with sexual conflict in *Drosophila melanogaster*. Behav. Genet 35:455–462. https://doi.org/10.1007/s10519-004-1246-8

Friberg U (2006) Male perception of female mating status: its effect on copulation duration, sperm defence and female fitness. Anim. Behav 72:1259–1268. https://doi.org/10.1016/j.anbehav.2006.03.021

Friberg U, Arnqvist G (2003) Fitness effects of female mate choice: preferred males are detrimental for *Drosophila melanogaster* females. J. Evol. Biol 16:797–811. https://doi.org/10.1046/j.1420-9101.2003.00597.x

García-Roa R, Chirinos V, Carazo P (2019) The ecology of sexual conflict: Temperature variation in the social environment can drastically modulate male harm to females. Funct. Ecol 33:681–692. https://besjournals.onlinelibrary.wiley.com/journal/13652435

Ghosh SM, Joshi A (2012) Evolution of reproductive isolation as a by-product of divergent life-history evolution in laboratory populations of *Drosophila melanogaster*. Ecol. Evol 2:3214–3226. https://doi.org/10.1002/ece3.413

Gomez-Llano MA, Bensch HM, Svensson EI (2018) Sexual conflict and ecology: species composition and male density interact to reduce male mating harassment and increase female survival. Evolution 72:906–915. https://doi.org/10.1111/evo.13457

Griffith SC (2019) Cooperation and coordination in socially monogamous birds: moving away from a focus on sexual conflict. Front. Ecol. Evol 7:455. https://doi.org/10.3389/fevo.2019.00455

Gromko MH (1987) Genetic constraint on the evolution of courtship behaviour in *Drosophila melanogaster*. Heredity 58:435–441. https://doi.org/10.1038/hdy.1987.72

Hartmann R, Loher W (1999) Post-mating effects in the grasshopper, *Gomphocerus rufus* L. mediated by the spermatheca. J. Comp. Physiol 184:325–332. https://doi.org/10.1007/s003590050330

Holland B, Rice WR (1998) Perspective: chase-away sexual selection: antagonistic seduction versus resistance. Evolution 52: 1–7. https://doi.org/10.1111/j.1558-5646.1998.tb05132.x

Holland B, Rice WR (1999) Experimental removal of sexual selection reverses intersexual antagonistic coevolution and removes a reproductive load. Proc. Natl. Acad. Sci. U.S.A. 96:5083–5088. http://dx.doi.org/10.1073/pnas.96.9.5083

Hooper AK, Spagopoulou F, Wylde Z, Maklakov AA, Bonduriansky R (2017) Ontogenetic timing as a condition-dependent life history trait: High-condition males develop quickly, peak early, and age fast. Evolution 71:671–685. https://doi.org/10.1111/evo.13172

Hunt J, Brooks R, Jennions MD, Smith MJ, Bentsen CL, Bussiere LF (2004) High-quality male field crickets invest heavily in sexual display but die young. Nature 432:1024–1027. https://doi.org/10.1038/nature03084

Jennions MD, Petrie M (2000) Why do females mate multiply? A review of the genetic benefits. Biol. Rev 75:21–64. https://doi.org/10.1017/S0006323199005423

Jiang PP, Bedhomme S, Prasad N, Chippindale A (2011) Sperm competition and mate harm unresponsive to male-limited selection in *Drosophila*: an evolving genetic architecture under domestication. Evol. int. j. org. evol 65:2448–2460. https://doi.org/10.1111/j.1558-5646.2011.01328.x

Johnstone RA, Keller L (2000) How males can gain by harming their mates: sexual conflict, seminal toxins, and the cost of mating. Am. Nat 156:368–377. https://doi.org/10.1086/303392

Kirkwood TB, Rose MR (1991) Evolution of senescence: late survival sacrificed for reproduction. Philos. Trans. R. Soc. Lond., B, Biol. Sci 332:15–24. https://doi.org/10.1098/rstb.1991.0028

Koene JM (2012) Sexual conflict in nonhuman animals. In: The Oxford handbook of sexual conflict in humans. http://dx.doi.org/10.1093/oxfordhb/9780195396706.013.0002

Kotiaho JS (2001) Costs of sexual traits: a mismatch between theoretical considerations and empirical evidence. Biol. Rev 76:365–376. https://doi.org/10.1017/s1464793101005711

Kotiaho JS, Simmons LW (2003) Longevity cost of reproduction for males but no longevity cost of mating or courtship for females in the male-dimorphic dung beetle Onthophagus binodis. J. Insect Physiol 49:817–822. https://doi.org/10.1016/S0022-1910(03)00117-3

Kuznetsova A, Brockhoff PB, Christensen RH (2017) lmerTest package: tests in linear mixed effects models. J. Stat. Softw 82:1–26. https://doi.org/10.18637/jss.v082.i13

Lane JE, Boutin S, Speakman JR, Humphries MM (2010) Energetic costs of male reproduction in a scramble competition mating system. J. Anim. Ecol 79:27–34. https://doi.org/10.1111/j.1365-2656.2009.01592.x

Le Galliard J-F, Fitze PS, Ferrière R, Clobert J (2005) Sex ratio bias, male aggression, and population collapse in lizards. Proc. Natl. Acad. Sci. U.S.A. 102:18231–18236. https://doi.org/10.1073/pnas.0505172102

Lemaître J-F, Ronget V, Tidière M, Allainé D, Berger V, Cohas A, Colchero F, Conde DA, Garratt M, Liker A (2020) Sex differences in adult lifespan and aging rates of mortality across wild mammals. Proc. Natl. Acad. Sci. U.S.A. 117:8546–8553. https://doi.org/10.1073/pnas.1911999117

Lenth R, Singmann H, Love J, Buerkner P, Herve M (2018) Emmeans: Estimated marginal means, aka least-squares means. R package version, 1, 3. https://doi.org/10.1080/00031305.1980.10483031

Linklater JR, Wertheim B, Wigby S, Chapman T (2007) Ejaculate depletion patterns evolve in response to experimental manipulation of sex ratio in *Drosophila melanogaster*. Evolution 61:2027–2034. https://doi.org/10.1111/j.1558-5646.2007.00157.x

Long T, Pischedda A, Nichols R, Rice W (2010) The timing of mating influences reproductive success in *Drosophila melanogaster:* implications for sexual conflict. J. Evol. Biol 23:1024–1032. https://doi.org/10.1111/j.1420-9101.2010.01973.x

MacPherson A, Yun L, Barrera TS, Agrawal AF, Rundle HD (2018) The effects of male harm vary with female quality and environmental complexity in *Drosophila melanogaster*. Biol. Lett 14:20180443. https://doi.org/10.1098/rsbl.2018.0443

Maklakov AA, Lummaa V (2013) Evolution of sex differences in lifespan and aging: causes and constraints. BioEssays 35:717–724. https://doi.org/10.1002/bies.201300021

Manier MK, Belote JM, Berben KS, Novikov D, Stuart WT, Pitnick S (2010) Resolving mechanisms of competitive fertilization success in *Drosophila melanogaster*. Science 328:354–357. https://doi.org/10.1126/science.1187096

Markow TA, Hanson SJ (1981) Multivariate analysis of *Drosophila* courtship. Proc. Natl. Acad. Sci. U.S.A. 78:430–434. https://doi.org/10.1073/pnas.78.1.430

Mital A, Sarangi M, Joshi A (2021) Evolution of lower levels of inter-locus sexual conflict in D. melanogaster populations under strong selection for rapid development. BioRxiv. https://doi.org/10.1101/2021.02.08.430125

Mital A, Sarangi M, Nandy B, Pandey N, Joshi A (2022) Shorter effective lifespan in laboratory populations of D. melanogaster might reduce sexual selection. Behav Ecol Sociobiol 76, 52. https://doi.org/10.1007/s00265-022-03158-w

Montrose VT, Harris WE, Moore P (2004) Sexual conflict and cooperation under naturally occurring male enforced monogamy. J. Evol. Biol 17:443–452. https://doi.org/10.1046/j.1420-9101.2003.00654.x

Morrow EH, Arnqvist G, Pitnick S (2003) Adaptation versus pleiotropy: why do males harm their mates? Behavioral Ecology 14:802–806. https://doi.org/10.1093/beheco/arg073

Nandy B, Chakraborty P, Gupta V, Ali SZ, Prasad NG (2013a) Sperm competitive ability evolves in response to experimental alteration of operational sex ratio. Evolution 67:2133–2141. https://doi.org/10.1111/evo.12076

Nandy B, Dasgupta P, Halder S, Verma T (2016) Plasticity in aggression and the correlated changes in the cost of reproduction in male Drosophila melanogaster. Anim. Behav 114:3–9. https://doi.org/10.1016/j.anbehav.2016.01.019

Nandy B, Gupta V, Sen S, Udaykumar N, Samant MA, Ali SZ, Prasad NG (2013b) Evolution of mate-harm, longevity and behaviour in male fruit flies subjected to different levels of inter locus conflict. BMC Evol. Biol 13:1–16. https://doi.org/10.1186/1471-2148-13-212

Nandy B, Gupta V, Udaykumar N, Samant M, Sen S, Prasad N (2014) Experimental evolution of female traits under different levels of intersexual conflict in *Drosophila melanogaster*. Evolution 68:412–425. https://doi.org/10.1111/evo.12271

Nandy B, Joshi A, Ali ZS, Sen S, Prasad NG (2012) Degree of adaptive male mate choice is positively correlated with female quality variance. Sci. Rep 2:1–8. https://doi:10.1038/srep00447

Nandy B, Prasad NG (2011) Reproductive behavior and fitness components in male Drosophila melaogaster are non-linearly affected by the number of male co-inhabitants early in adult life. J. Insect Sci 11:67. https://doi.org/10.1673/031.011.6701

Parker GA (1979) Sexual selection and sexual conflict. Sexual selection and reproductive competition in insects 123:166. https://doi.org/10.1016/B978-0-12-108750-0.50010-0

Parker GA (2006) Sexual conflict over mating and fertilization: an overview. Philos. Trans. R. Soc. Lond., B, Biol. Sci 361:235–259. https://dx.doi.org/10.1098%2Frstb.2005.1785

Partridge L, Farquhar M (1981) Sexual activity reduces lifespan of male fruit flies. Nature 294:580–582. https://doi.org/10.1038/294580a0

Passananti HB, Matos M (2004) Methuselah flies: a case study in the evolution of aging. World Scientific. https://doi.org/10.1086/433090

Pitnick S (1991) Male size influences mate fecundity and remating interval in Drosophila melanogaster. Anim. Behav 41:735–45. https://doi.org/10.1016/S0003-3472(05)80340-9

Pitnick S, García–González F (2002) Harm to females increases with male body size in *Drosophila melanogaster*. Proc. Royal Soc 269:1821–1828. https://doi.org/10.1098/rspb.2002.2090

Pitnick S, Miller GT, Reagan J, Holland B (2001) Males’ evolutionary responses to experimental removal of sexual selection. Proc. Royal Soc 268:1071–1080. https://doi.org/10.1098/rspb.2001.1621

Poiani A (2006) Complexity of seminal fluid: a review. Behav. Ecol. Sociobiol 60:289–310. https://doi.org/10.1007/s00265-006-0178-0

Promislow DE (1992) Costs of sexual selection in natural populations of mammals. Proc. Royal Soc 247:203–210. https://doi.org/10.1098/rspb.1992.0030

Prowse N, Partridge L (1997) The effects of reproduction on longevity and fertility in male *Drosophila melanogaster*. J. Insect Physiol 43:501–512. https://doi.org/10.1016/S0022-1910(97)00014-0

Queller DC, Strassmann JE (2018) Evolutionary conflict. Annu Rev Ecol Evol Syst 49:73–93. https://doi.org/10.1146/annurev-ecolsys-110617-062527

Rankin DJ, Bargum K, Kokko H (2007) The tragedy of the commons in evolutionary biology. Trends Ecol. Evol 22:643–651. https://doi.org/10.1016/j.tree.2007.07.009

Rankin DJ, Dieckmann U, Kokko H (2011) Sexual conflict and the tragedy of the commons. Am. Nat 177:780–791. https://doi.org/10.1086/659947

Rankin DJ, Kokko H (2006) Sex, death and tragedy. Trends Ecol. Evol 21:225–226. https://doi.org/10.1016/j.tree.2006.02.013

Rice WR (1992) Sexually antagonistic genes: experimental evidence. Science 256:1436–1439. http://www.jstor.org/stable/2876947

Rice WR (1996) Sexually antagonistic male adaptation triggered by experimental arrest of female evolution. Nature 381:232–234. https://doi.org/10.1038/381232a0

Ritchie M, Gleason J (1995) Rapid evolution of courtship song pattern in *Drosophila willistoni* sibling species. J. Evol. Biol 8:463–479. https://doi.org/10.1046/j.1420-9101.1995.8040463.x

Rönn J, Katvala M, Arnqvist G (2007) Coevolution between harmful male genitalia and female resistance in seed beetles. Proc. Natl. Acad. Sci. U.S.A. 104:10921–10925. https://doi.org/10.1073/pnas.0701170104

Rose MR, Charlesworth B (1981) Genetics of life history in *Drosophila melanogaster*. II. Exploratory selection experiments. Genetics 97:187–196. https://doi.org/10.1093/genetics/97.1.187

Rose MR, Vu LN, Park SU, Graves Jr JL (1992) Selection on stress resistance increases longevity in *Drosophila melanogaster*. Exp. Gerontol 27:241–250. https://doi.org/10.1016/0531-5565(92)90048-5

Rostant WG, Mason JS, de Coriolis JC, Chapman T (2020) Resource-dependent evolution of female resistance responses to sexual conflict. Evol. Lett 4:54–64. https://doi.org/10.1002/evl3.153

Ruedi EA, Hughes KA (2008) Natural genetic variation in complex mating behaviors of male *Drosophila melanogaster*. Behav. Genet 38:424–436. https://doi.org/10.1007/s10519-008-9204-5

Service PM (1993) Laboratory evolution of longevity and reproductive fitness components in male fruit flies: mating ability. Evolution 47:387–399. https://doi.org/10.2307/2410059

Singh SR, Singh BN (2004) Female remating in *Drosophila:* comparison of duration of copulation between first and second matings in six species. Curr. Sci 86:465–470. https://www.jstor.org/stable/24108745

Snook RR (2001) Sexual selection: conflict, kindness and chicanery. Curr. Biol 11:R337–R341. https://doi.org/10.1016/S0960-9822(01)00188-9

Snook R, Robertson A, Crudgington H, Ritchie M (2005) Experimental manipulation of sexual selection and the evolution of courtship song in *Drosophila pseudoobscura*. Behavior genetics 35:245–255. https://doi.org/10.1007/s10519-005-3217-0

Snow SS, Alonzo SH, Servedio MR, Prum RO (2019) Female resistance to sexual coercion can evolve to pre-serve the indirect benefits of mate choice. J. Evol. Biol 32:545–558. https://doi.org/10.1111/jeb.13436

Sokal RR, Rohlf FJ (1995) Biometry. New York: W. H. H Freeman and Company. http://dx.doi.org/10.2307/2343822

Stearns SC (1989) Trade-offs in life-history evolution. Funct. Ecol 3:259–268. https://doi.org/10.2307/2389364

Stewart AD, Morrow EH, and Rice WR (2005) Assessing putative interlocus sexual conflict in Drosophila mel-anogaster using experimental evolution. Proc. Royal Soc. B: Biol. Sci 272: 2029–2035. https://doi.org/10.1098/rspb.2005.3182

Tatar M, Carey JR, Vaupel JW (1993) Long-term cost of reproduction with and without accelerated senescence in *Callosobruchus maculatus:* analysis of age-specific mortality. Evolution 47:1302–1312. https://doi.org/10.1111/j.1558-5646.1993.tb02156.x

Von Schilcher F, Dow M (1977) Courtship behaviour in Drosophila: sexual isolation or sexual selection? Z Tierpsychol 43:304–310. https://doi.org/10.1111/j.1439-0310.1977.tb00077.x

Wedell N, Kvarnemo C, Tregenza T (2006) Sexual conflict and life histories. Anim. Behav 71:999–1011. https://doi.org/10.1016/j.anbehav.2005.06.023

Wigby S, Brown NC, Allen SE, Misra S, Sitnik JL, Sepil I, Clark AG, Wolfner MF (2020) The *Drosophila* seminal proteome and its role in postcopulatory sexual selection. Philos. Trans. R. Soc. Lond., B, Biol. Sci 375:20200072. https://doi.org/10.1098/rstb.2020.0072

Wigby S, Chapman T (2004) Female resistance to male harm evolves in response to manipulation of sexual conflict. Evolution 58:1028–1037. https://doi.org/10.1111/j.0014-3820.2004.tb00436.x

Wigby S, Chapman T (2005) Sex peptide causes mating costs in female *Drosophila melanogaster*. Curr. Biol 15:316–321. https://doi.org/10.1016/j.cub.2005.01.051

Wolfner M (2002) The gifts that keep on giving: physiological functions and evolutionary dynamics of male seminal proteins in *Drosophila*. Heredity 88:85–93. https://doi.org/10.1038/sj.hdy.6800017

Wolfner MF (1997) Tokens of love: functions and regulation of *Drosophila* male accessory gland products. Insect Biochem. Mol. Biol 27:179–192. https://doi.org/10.1016/S0965-1748(96)00084-7

Yun L, Chen PJ, Singh A, Agrawal AF, Rundle HD (2017) The physical environment mediates male harm and its effect on selection in females. Proc. Royal Soc. B 284:20170424. https://doi.org/10.1098/rspb.2017.0424

Zar JH (1999) Biostatistical analysis. Pearson Education India.

