## Supplementary material for "Evolution of reduced mate harming tendency of males in *Drosophila melanogaster* populations selected for faster life history": Supplemnetary information

**Supplementary information:**

**A. Experimental populations:**

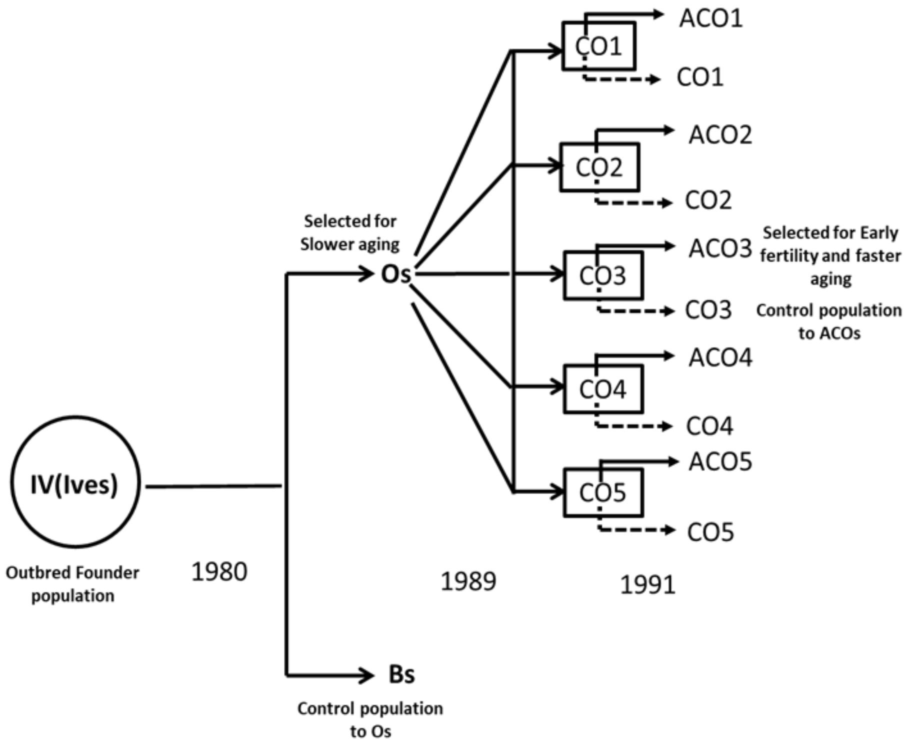


**Fig. S1** History of the experimental populations including phylogenetic relationships. Five replicates of IV population were derived from wild-caught flies in 1975, and were maintained as outbred population. IV populations were maintained on 2-week discrete generation cycle. Subsequently, five replicates of the baseline B-populations were established ($B_{1-5}$), which were maintained under a regime identic to that of the IV populations. Five replicates of the O populations ($O_{1-5}$), along with their matched controls ($CO_{1-5}$), selected for postponed senescence were derived in 1980. In 1991, CO populations were used to establish five replicates of the ACO populations ($ACO_{1-5}$), which were subjected to selection for accelerated development and early reproduction

**B. Dry whole body weight of experimental males at eclosion:**

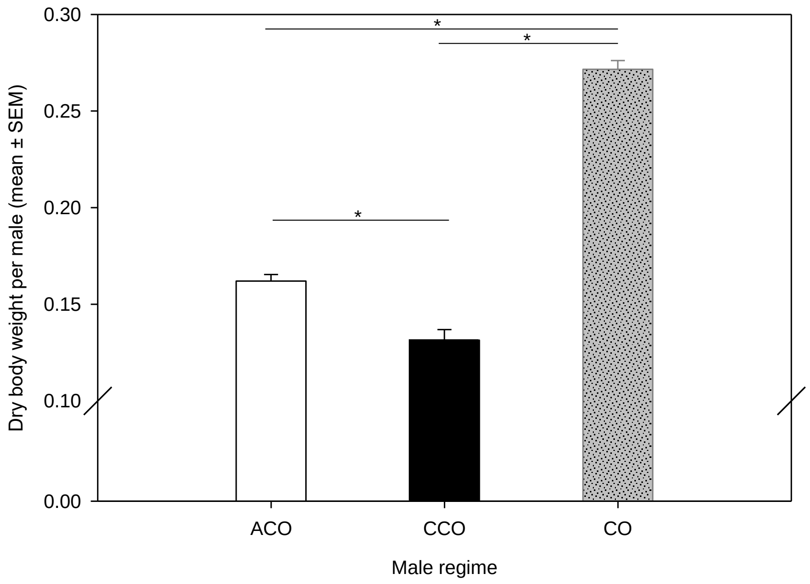


**Fig. S2** Effect of male regime on mean dry body weight at eclosion for experimental males in the main assay. Means were calculated over five blocks of the entire experiment. The vertical bars indicate the mean across all replicate populations. Error bars represent the standard errors of means (SEM). Significant differences are marked with horizontal line and an asterix (*), determined using Tukey's HSD

| Block | SS | DF | MS | F | p |
|---|---|---|---|---|---|
| **1** | 0.103 | 2 | 0.051 | 67.653 | **<0.001** |
| **2** | 0.107 | 2 | 0.054 | 179.436 | **<0.001** |
| **3** | 0.193 | 2 | 0.097 | 95.736 | **<0.001** |
| **4** | 0.0852 | 2 | 0.043 | 400.086 | **<0.001** |
| **5** | 0.087 | 2 | 0.044 | 49.853 | **<0.001** |

**Table S1** Summary of the results of one-way ANOVA on dry body weight data from individual blocks using male regime as categorical predictor. Dry body weight was measured in group of five flies which was then used to calculate per male body weight and this value was used as unit of analysis. All tests were done considering α=0.05 and significant p-values are mentioned in bold font style

**C. Additional analyses on the age specific per-capita fecundity:**

To analyse the age specific per capita fecundity, the following model was used for the linear mixed-effect model in R version 3.6.1 using `lme4` package (R Development Core Team, 2019) and `lmerTest`:

PCF ~ MALE REGIME+ AGE+ MALE REGIME:AGE + (1|BLOCK/VIAL_ID) + (1|BLOCK:MALE REGIME) + (1|BLOCK:AGE) + (1|BLOCK:AGE:MALE REGIME)

| Effect | SS | DF | MS | Den DF | F | p |
|---|---|---|---|---|---|---|
| Male regime | 234.644 | 2 | 117.322 | 7.995 | 16.0997 | **0.001** |
| Age | 218.392 | 4 | 54.598 | 16.006 | 7.4923 | **0.001** |
| Male regime × Age | 45.664 | 8 | 5.708 | 31.936 | 0.7833 | 0.620 |

**Table S2** Summary of results of linear mixed-effect model analysis (LMM) on per capita fecundity under CE male exposure type. Male regime and age were fitted as categorical fixed effect and block as a random factor in per capita fecundity analysis. All tests were done considering α=0.05 and significant p-values are mentioned in bold font style

| Effect | npar | AIC | LogLik | Chisq | DF | Pr(>Chisq) |
|---|---|---|---|---|---|---|
| Block × Male regime | 20 | 3810.500 | -1885.2 | 0.621 | 1 | 0.430 |
| Block × Age | 20 | 3828.000 | -1894.0 | 18.160 | 1 | **<0.001** |
| Block × Male regime × age | 20 | 3828.200 | -1894.1 | 18.339 | 1 | **<0.001** |
| Block | 19 | 3850.600 | -1906.3 | 42.760 | 2 | **<0.001** |

**Table S3** Effect of block as a random factor and those of all two-way interactions involving block on age specific per capita fecundity. This was done using a linear mixed-effect model in R version 3.6.1 using `lme4` package (R Development Core Team, 2019) and `lmerTest`. Significant effects (p < 0.05) are shown in bold font style

<u>Block-wise analysis:</u>

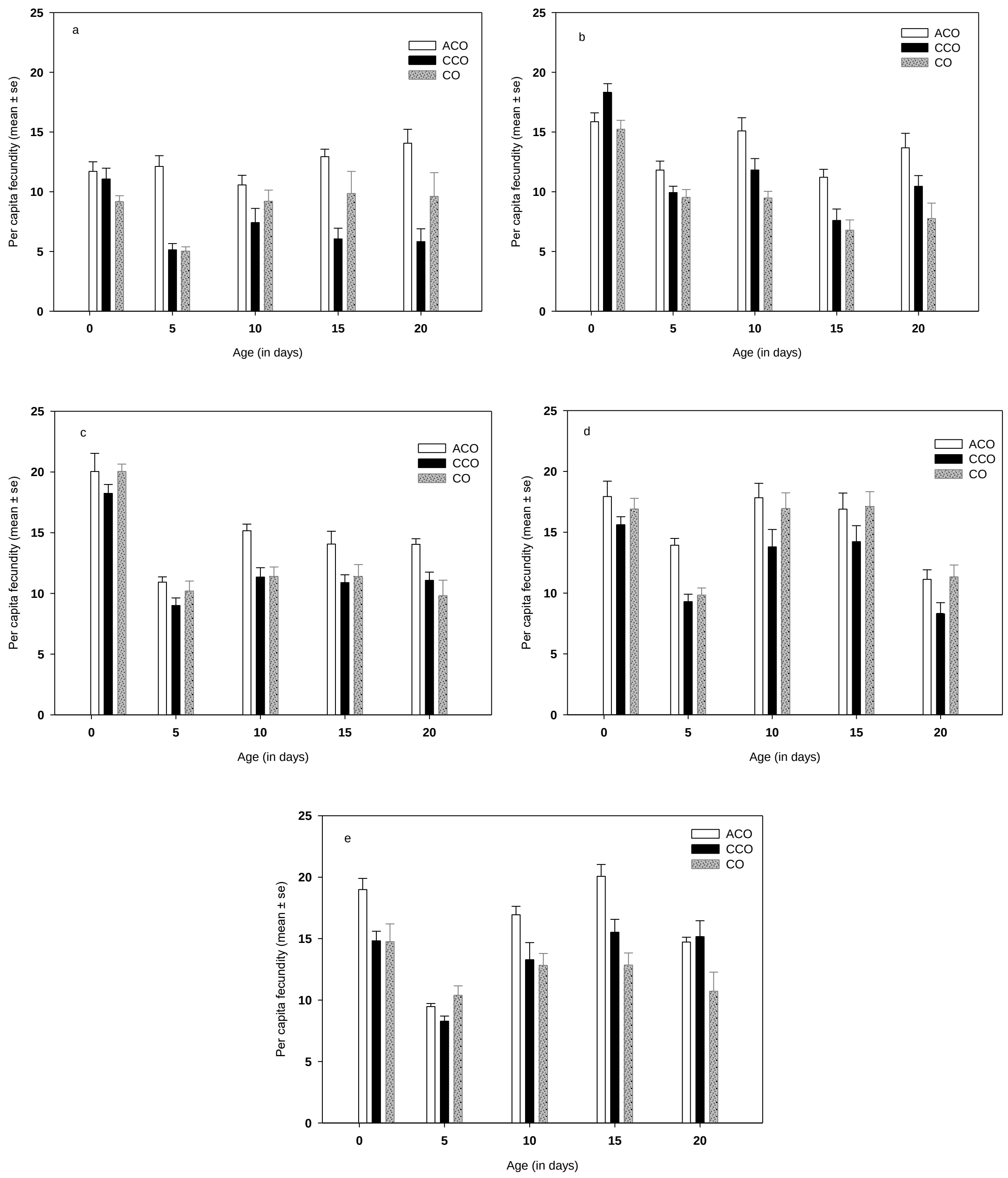


**Fig. S3** Effect of exposure to treatment males (ACO/CCO/CO) on age specific per capita fecundity of experimental Oregon R females. The panels show results from (a) Block 1, (b) Block 2, (c) Block 3, (d) Block 4, and (e) Block 5.

| Effect | Block | SS | MS | Num DF | Den DF | F | p |
|---|---|---|---|---|---|---|---|
| Male regime (MR) | **1** | 404.01 | 202.003 | 2 | 26.206 | 22.457 | **<0.001** |
| Age | | 164.91 | 41.228 | 4 | 102.750 | 4.583 | **0.002** |
| MR × Age | | 266.03 | 33.254 | 8 | 102.747 | 3.697 | **0.001** |
| Male regime (MR) | **2** | 115.95 | 57.973 | 2 | 27 | 10.756 | **<0.001** |
| Age | | 1071.24 | 267.811 | 4 | 108 | 49.687 | **<0.001** |
| MR × Age | | 172.28 | 21.535 | 8 | 108 | 3.995 | **<0.001** |
| Male regime (MR) | **3** | 103.77 | 51.89 | 2 | 26 | 9.204 | **<0.001** |
| Age | | 1528.12 | 382.03 | 4 | 104 | 67.771 | **<0.001** |
| MR × Age | | 69.72 | 8.71 | 8 | 104 | 1.546 | 0.150 |
| Male regime (MR) | **4** | 169.58 | 84.790 | 2 | 27 | 8.8503 | **0.001** |
| Age | | 1200.77 | 300.193 | 4 | 104 | 31.334 | **<0.001** |
| MR × Age | | 73.43 | 9.178 | 8 | 104 | 0.9580 | 0.473 |
| Male regime (MR) | **5** | 121.32 | 60.661 | 2 | 26.805 | 8.8366 | **0.001** |
| Age | | 931.38 | 232.846 | 4 | 105.356 | 33.9192 | **<0.001** |
| MR × Age | | 250.10 | 31.262 | 8 | 105.339 | 4.5541 | **<0.001** |

**Table S4** Summary of results of linear mixed-effect model (LMM) on female per capita fecundity under CE exposure type for each block separately. Male regime, age and their two-way interaction were modelled as fixed factors. All tests were done considering α=0.05 and significant p-values are mentioned in bold font style

**D. Block-wise analyses on the cumulative female mortality:**

| Effects | Block | SS | DF | MS | F | p |
|---|---|---|---|---|---|---|
| Male regime (MR) | **1** | 0.343 | 2 | 0.171 | 10.188 | **<0.001** |
| Male exposure type (MET) | | 0.620 | 1 | 0.620 | 36.842 | **<0.001** |
| MR × MET | | 0.500 | 2 | 0.250 | 14.861 | **<0.001** |
| Male regime (MR) | **2** | 0.822 | 2 | 0.411 | 18.470 | **<0.001** |
| Exposure type (MET) | | 1.414 | 1 | 1.414 | 63.569 | **<0.001** |
| MR × MET | | 1.041 | 2 | 0.520 | 23.395 | **<0.001** |
| Male regime (MR) | **3** | 0.021 | 2 | 0.010 | 0.619 | 0.542 |
| Exposure type (MET) | | 0.370 | 1 | 0.370 | 22.005 | **<0.001** |
| MR × MET | | 0.120 | 2 | 0.060 | 3.572 | **0.035** |
| Male regime (MR) | **4** | 0.732 | 2 | 0.366 | 17.754 | **<0.001** |
| Exposure type (MET) | | 0.677 | 1 | 0.677 | 32.851 | **<0.001** |
| MR × MET | | 0.747 | 2 | 0.373 | 18.112 | **<0.001** |
| Male regime (MR) | **5** | 0.357 | 2 | 0.178 | 10.645 | **<0.001** |
| Exposure type (MET) | | 0.344 | 1 | 0.344 | 20.548 | **<0.001** |
| MR × MET | | 0.387 | 2 | 0.194 | 11.569 | **<0.001** |

**Table S5** Summary of results of three-factor ANOVA on cumulative female mortality done for each block separately. Male regime and male-exposure type were modelled as a fixed factor in female mortality. All tests were done considering $\alpha$=0.05 and significant p-values are mentioned in bold font style

**E. Additional analysis age specific courtship frequency:**

| Effect | npar | AIC | LogLik | Chisq | DF | Pr(>Chisq) |
|---|---|---|---|---|---|---|
| Block × Male regime | 15 | 210.980 | -90.491 | 0.905 | 1 | 0.341 |
| Block × Age | 15 | 212.640 | -91.318 | 2.559 | 1 | 0.110 |
| Block × Male regime × age | 15 | 210.710 | -90.353 | 0.6283 | 1 | 0.428 |
| Block | 14 | 208.250 | -90.122 | 0.168 | 2 | 0.919 |

**Table S6** Effect of block as a random factor and those of all two-way interactions involving block on age specific courtship frequency. This was done using a linear mixed-effect model in R version 3.6.1 using `lme4` package (R Development Core Team, 2019) and `lmerTest`. Significant effects ($p < 0.05$) are shown in bold font style

**F. Copulation duration assay:**

To compare the mating behaviour of the ACO and CO regime, mating behaviour assays were set up using experimental flies generated from standardized populations. Five virgin males and five virgin females from the same population were put together in fresh food vials to set up the mating behaviour assay. Ten replicate vials were set up for each population. This assay was performed on three of the five blocks – $ACO_1/CO_1$, $ACO_2/CO_2$, $ACO_3/CO_3$. Following the combination of the sexes, these vials were checked every 2 minutes, and the total number of copulating pairs were counted until all copulating pairs have completed mating (i.e., disengaged from mounted position). Mean copulation duration (CD) for a vial was calculated following an algorithm mentioned below, where N is the total number of females, $n_x$ is count of copulating pair at $x^{th}$ minutes since the onset of the mating trial.

$$CD = \frac{\sum(n_{x-2} - n_x)x}{N} - ML \qquad \text{(For all values of x, for which } n_{x-2} > n_x\text{)}$$

In the above algorithm, ML (Mating latency: time required by virgin pairs to start mating following combination of the sexes) was calculated using following algorithm, where N is the total number of females, $n_y$ is count of copulating pair at $y^{th}$ minutes since the onset of the mating trial.

$$ML = \frac{\sum(n_y - n_{y-2})y}{N} \qquad \text{(For all values of y, for which } n_{y-2} > n_y\text{)}$$

CD data was analysed using two factor mixed model ANOVA using selection regime as a fixed factor and block as a random factor. The analysis revealed a significant effect of selection regime, with CD being significantly shorter in the ACOs. On an average, we found a 20% reduction in CD for the ACOs.

| Effect | SS | DF | MS | Den DF | F | p |
|---|---|---|---|---|---|---|
| Selection regime | 260.970 | 1 | 260.970 | 2.000 | 102.886 | **0.009** |
| Block | 12.200 | 2 | 6.100 | 2.000 | 2.405 | 0.294 |
| Selection regime × Block | 5.070 | 2 | 2.540 | 54.000 | 0.197 | 0.822 |

**Table S7** Results of the two-factor ANOVA on mean copulation duration, CD across ACO and CO regime. Selection regime was modelled as a fixed factor and block as a random factor in copulation duration analysis.

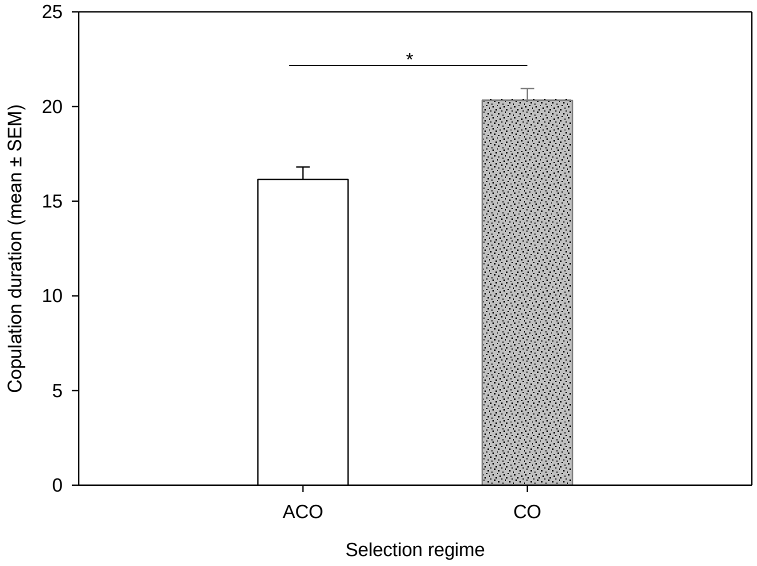


**Fig. S4** Mean copulation duration across ACO and CO males. Mean CD was calculated for five males in a vial following the algorithm given above. The values obtained was then used as unit of analysis. The vertical bars indicate the mean across all replicate populations. Error bars represent the standard errors of means (SEM). Significant differences are marked with horizontal line and an asterix (*), determined using Tukey's HSD

**G. Remating frequency assay:**

To assess the scope of sexual selection, especially post-copulatory sexual selection, we quantified how often females in ACO populations undergo re-mating within the 24 hours of their usual adult life. For comparison, we also quantified the same for CO populations. This was done for three of the five replicate populations ($ACO_2/CO_2$, $ACO_3/CO_3$ and $ACO_5/CO_5$). Since, the purpose of this assay was to get a measure of the extent to which females in these populations undergo remating, we did not use standardized flies. Experimental flies were generated by collecting eggs directly from the running stock populations, without passing a subset through one generation of common gardening maintenance. Otherwise, the method of collection of eggs, larval density in the growth vials, and all other conditions were identical to those of the main experiment (mentioned in the Methods section). Flies were collected as virgin under light $CO_2$ anaesthesia, and both males and females were housed in single-sex vials (10 individuals per vial) until the observation. When the adults were 1-2 day old, 35 mating vials were set up for each population. These mating vials were set up by combining one vial of each sex, so that 10 virgin males and 10 virgin females from the same population were held in these vials. The vials were manually observed for 1-2 hours to ensure that all females went through a single round of mating. Afterwards, flies from all 35 vials (for a population) were transferred to a transparent cage (1.5-liter volume). Each cage had ~700 flies (i.e., 350 males, 350 females). The cages were left undisturbed for ~30 minutes for flies to acclimatize before starting the observation. For the next 24 hours, the number of copulating pairs in a cage was counted every 15 minutes. Since, all females completed their first mating before being introduced in the observation cage, all copulating pairs observed in the cage were considered remating events. This count was considered a measure of remating frequency. Mean remating frequency in 24 hours across two selection regime is plotted below. Mean remating frequency for ACO regime was found to be, 3.64±0.206 (mean±sem) and for CO regime 2.91±0.246 (mean±sem).

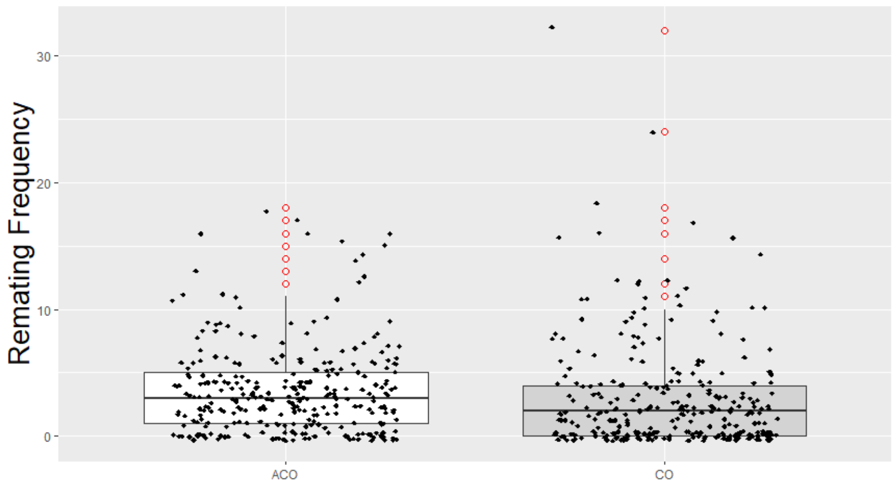


**Fig. S5** Box plot for mean remating frequency across ACO and CO selection regime. Box plot represents median and interquartile range. Red circles represent outliers. Effect of male regime on re-mating frequency was not found to be significant
